# TLR7-Dependent Autoimmunity Remodels Intestinal Immune Cell Dissemination

**DOI:** 10.64898/2026.10.05.756804

**Authors:** Katherine S. Saldana-Morales, Miki Kume, Silvia R. Davila-Paico, Maricielo Vaella-Alarcon, Olivia Bache, Daniel F. Zegarra-Ruiz

## Abstract

Intestinal immune cell dissemination has emerged as a fundamental mechanism of tissue-to-tissue immune communication that shapes systemic immunity, yet how this physiological process changes across different stages of systemic autoimmunity remains poorly understood. Here, using photoconversion, inducible and spontaneous models of TLR7-dependent autoimmunity, and microbiota perturbation, we define how autoimmunity reshapes intestinal immune dissemination. TLR7-dependent autoimmunity progressively suppresses dissemination in a tissue-dependent manner across gut-associated and systemic tissues while selectively reconfiguring the intestinal emigrant pool, characterized by reduced lymphocyte dissemination and increased representation of myeloid populations, particularly dendritic cells. This divergence is already evident in young TLR7-transgenic mice before detectable splenomegaly. Moreover, microbial regulation of intestinal dissemination depends on timing and inflammatory context, with antibiotic treatment during ongoing autoimmunity producing broader changes in intestinal immune dissemination. Together, our findings establish remodeling of physiological intestinal immune dissemination as an early feature of TLR7-dependent autoimmunity and reveal how chronic inflammation reshapes tissue-to-tissue immune communication.

## INTRODUCTION

Immune cells continuously disseminate from the intestine to peripheral lymphoid and non-lymphoid tissues as part of normal immune homeostasis. Previous intestinal migration studies revealed unexpectedly broad movement of both lymphoid and myeloid populations from the gut at steady state, and more recent systematic analyses have shown that individual intestinal immune populations follow distinct migratory trajectories to distant organs ^1,2^. Importantly, this trafficking can transmit information from the intestinal environment beyond the mucosa. In early life, microbiota-influenced plasmacytoid dendritic cells migrate from the colon to the thymus and regulate the development of PLZF⁺ innate lymphocytes ^3^. Gut-derived dendritic cells can also transport microbiota-derived antigens to the thymus, where they shape the development of microbiota-specific T cells ^4^. Together, these studies establish physiological intestinal immune dissemination as a mechanism through which the gut communicates with distant immune tissues and contributes to systemic immune homeostasis.

Migration of immune cells from the gut is sensitive to changes in the intestinal environment and can be reshaped during disease ^2,5–8^. Microbiota depletion reduces the traffic of gut-derived immune cells, and intestinal inflammation similarly suppresses dissemination of many populations, with dendritic cells representing a notable exception during inflammation ^2^. Migration of immune cells from the gut to systemic tissues can also directly contribute to extraintestinal pathology. In autoimmune arthritis, segmented filamentous bacteria promote the differentiation and systemic egress of Peyer’s patch T follicular helper cells that amplify autoantibody responses and disease ^6^, while gut-derived Th17 cells can acquire a highly migratory Tfh/17 phenotype and disseminate to systemic sites where they promote germinal center responses ^5^. Together, these studies demonstrate that intestinal immune dissemination is dynamically regulated by microbial and inflammatory signals and can transmit pathogenic immune responses beyond the gut. However, they have largely defined individual migratory populations or settings of acute intestinal inflammation, leaving unresolved the question of whether chronic systemic autoimmunity progressively remodels the physiological dissemination program as a whole.

Systemic lupus erythematosus provides a particularly relevant setting for examining how chronic inflammation reshapes inter-organ immune trafficking. Lupus is characterized by persistent inflammation and altered chemokine signaling ^9–11^, together with intestinal dysbiosis and altered host responses to the microbiota ^12–16^, all of which could influence immune cell movement between tissues. Aberrant Toll-like receptor 7 (TLR7) signaling is a major driver of lupus pathogenesis, and TLR7-dependent lupus models exhibit microbiota-dependent disease in which intestinal pathobionts can amplify systemic inflammation, making this a tractable context for studying these changes across disease progression ^16–19^. Consistent with the ability of lupus-associated inflammation to redirect immune cell trafficking, inflammatory cells originating in the skin can migrate to draining lymph nodes or disseminate to distant organs, including the kidney ^20,21^. However, whether chronic TLR7-dependent autoimmunity similarly remodels the physiological dissemination of immune cells from the intestine remains unknown.

Here, using intestinal photoconversion, inducible and spontaneous models of TLR7-dependent autoimmunity, and microbiota perturbation, we define how physiological intestinal immune dissemination is reorganized across autoimmune progression. We show that TLR7-dependent autoimmunity progressively suppresses intestinal immune dissemination while selectively shifting the emigrant pool from predominantly lymphoid populations toward increased representation of myeloid cells, particularly dendritic cells. This reorganization is already evident before established systemic pathology. Moreover, the influence of the intestinal microbiota on immune dissemination depends on inflammatory context and timing, becoming broader during systemic autoimmunity. Together, these findings establish remodeling of physiological intestinal immune dissemination as an early and dynamic feature of TLR7-dependent autoimmunity.

## RESULTS

### TLR7-dependent autoimmunity progressively suppresses physiological intestinal immune dissemination

Chemokine dysregulation and altered immune cell recruitment are prominent features of systemic autoimmunity, but whether intestinal trafficking programs are similarly remodeled in TLR7-dependent disease remains unknown ^9,22,23^. We therefore examined ileal transcriptional profiles from B6 mice treated with topical imiquimod (IMQ), a TLR7 agonist that induces progressive systemic autoimmunity ^16,18^, and from TLR7.1 transgenic (TLR7.1 Tg) mice, which spontaneously develop TLR7-dependent systemic autoimmune disease ^16,17^. Across both models, transcriptional profiling revealed broad remodeling of multiple chemokine ligands and receptors (Fig. S1A), indicating substantial reorganization of intestinal programs controlling immune cell movement.

These transcriptional changes raised the question of whether intestinal trafficking programs were accompanied by changes in the physiological dissemination of immune cells from the intestine. To directly examine this process in vivo, we used KikGR photoconvertible mice, which allow immune cells originating from a defined intestinal site to be tracked after photoconversion ^24^. We selectively exposed the cecum to 405-nm light, converting KikGR-expressing cells within the targeted region from GFP to RFP, and refer to these photoconverted RFP⁺ cells as gut-derived cells throughout the study. We then assessed their appearance in distant tissues 24 hours later (Fig. S1B). Photoconversion efficiently labeled leukocytes within the cecum while producing negligible RFP signal in the mesenteric lymph nodes (MLN), blood, spleen, or ileum immediately after conversion. In contrast, RFP⁺ cells were readily detected in these tissues 24 hours later, demonstrating dissemination from the photoconverted intestinal region rather than nonspecific or leaky photoconversion (Fig. S1C). Intravenous CD45 labeling further showed that, although circulating blood leukocytes were uniformly labeled, fewer than 1% of RFP⁺ cells recovered from the photoconverted cecum or spleen were intravascular, excluding blood contamination as a major source of the migratory population (Fig. S1D). Together, these controls established localized cecal photoconversion as a spatially restricted approach for tracking physiological dissemination of gut-derived immune cells in vivo.

Having established this system, we next applied it to inducible and spontaneous TLR7-dependent autoimmunity (Fig. 1A). KikGR mice beginning at approximately 7 weeks of age were treated chronically with topical IMQ and analyzed after 2, 4, and 6 weeks of treatment. In parallel, we generated KikGR × TLR7.1 Tg mice (KT mice) to combine spontaneous TLR7-dependent autoimmunity with in vivo tracking of intestinal emigrants. Adult KT mice were analyzed at approximately 12 weeks of age and compared with age-matched healthy littermate controls, placing them near the terminal time point of the IMQ cohort. By the latest IMQ time point, IMQ-treated mice and adult KT mice exhibited similar levels of anti-ssRNA and anti-dsDNA autoantibodies, whereas KT mice developed substantially greater splenic enlargement, consistent with more advanced systemic pathology (Fig. S1E, F). Together, these models allowed us to examine intestinal immune dissemination across increasing disease severity, from progressive IMQ-induced autoimmunity to the more advanced systemic pathology of adult KT mice.

**Figure 1.**
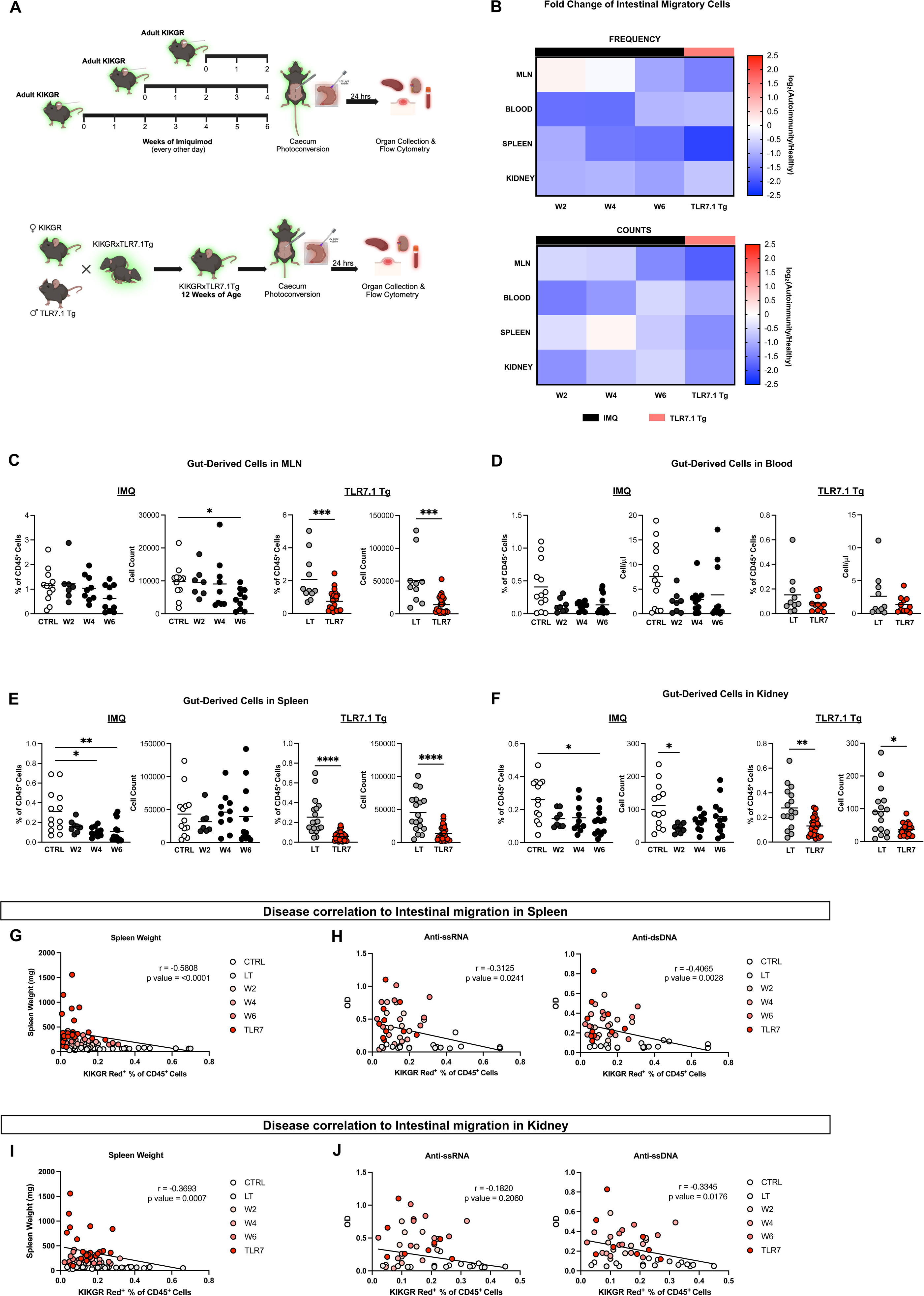
TLR7-dependent autoimmunity progressively suppresses physiological intestinal immune dissemination. (A) Experimental design for photoconversion in IMQ-induced model and TLR7.1Tg model. For the IMQ model, mice were treated topically with IMQ 3 times a week and underwent photoconversion at either 2 weeks, 4 weeks, or 6 weeks of treatment. For a TLR7.1Tg, we crossed KikGR mice with TLR7.1Tg to obtain KikGRxTLR7.1Tg; these mice underwent photoconversion at 12 weeks of age. In all cases, euthanasia and tissue collection were performed 24 hours after photoconversion. (B) Heatmaps showing fold change frequency and absolute counts of intestinal migratory cells (RFP^+^) in the MLN, blood, spleen and kidney. Values represent IMQ-treated KikGR mice at weeks 2, 4 and 6 (W2, W4, W6) relative to healthy KikGR controls (CTRL), and in KT adult mice (TLR7.1Tg) relative to their healthy littermate controls (LT). (C) Gut-derived RFP^+^ cell frequency and absolute counts in IMQ-treated KikGR mice at weeks 2, 4 and 6 and in KT adult mice (TLR7.1Tg) in the MLN (D) blood (E) spleen and (F) Kidney. (C-F, n = 6-27 per group). (G) Correlation between gut-derived cells frequency (KikGR Red^+^ % of CD45^+^ cells) in the spleen and spleen weights, in IMQ-treated KikGR mice at weeks 2, 4 and 6 and in KT adult mice (TLR7.1Tg) (n = 8-23 per group). (H) Correlation between gut-derived cells frequency (KikGR Red^+^ % of CD45^+^ cells) in the spleen and anti-ssRNA and anti-dsDNA autoantibody OD values, in IMQ-treated KikGR mice at weeks 2, 4 and 6 and in KT adult mice (TLR7.1Tg) (n = 8-10 per group). (I) Correlation between gut-derived cell frequency (KikGR Red^+^ % of CD45^+^ cells) in the Kidney and spleen weights, in IMQ-treated KikGR mice at weeks 2, 4 and 6 and in KT adult mice (TLR7.1Tg) (n = 8-23). (J) Correlation between gut-derived cell frequency (KikGR Red^+^ % of CD45^+^ cells) in the kidney and anti-ssRNA and anti-dsDNA autoantibody OD values, in IMQ-treated KikGR mice at weeks 2, 4 and 6 and in KT adult mice (TLR7.1Tg) (n = 8-10 per group). Data are shown as means. Statistical significance was determined by Kruskal-Wallis test with Dunn’s multiple comparison test (for more than two groups) and unpaired two-tailed Mann-Whitney U test (for two groups). Correlations were assessed using Spearman’s rank correlation. *\*p < 0.05, **p < 0.01, ***p < 0.001*.

Using this validated approach, we found that TLR7-dependent autoimmunity suppressed intestinal immune dissemination in a tissue-dependent manner. During IMQ-induced disease, gut-derived RFP⁺ cells progressively decreased in several peripheral compartments, with reduced cell numbers in the MLN and reduced frequencies in the spleen and kidney at later stages of disease, whereas dissemination through the blood remained comparatively preserved (Fig. 1B-F). The phenotype was more pronounced in adult KT mice, in which both the frequency and number of gut-derived RFP⁺ cells were markedly reduced in the MLN, spleen, and kidney, while blood dissemination again showed comparatively limited changes (Fig. 1B-F). Consistent with this progression, the frequency of gut-derived RFP⁺ cells in the spleen correlated inversely with splenic enlargement and anti-ssRNA and anti-dsDNA autoantibody levels, while kidney dissemination correlated inversely with splenic enlargement and anti-dsDNA autoantibodies (Fig. 1G-J). Thus, increasing severity of TLR7-dependent autoimmunity was associated with broader suppression of physiological intestinal immune dissemination across gut-associated and systemic tissues. We next asked whether systemic autoimmunity similarly reduced the dissemination of all intestinal immune cell populations or instead selectively reconfigured the composition of intestinal emigrants.

### Systemic autoimmunity selectively reconfigures the cellular composition of intestinal emigrants

The reduction in total gut-derived RFP⁺ cells raised the question of whether TLR7-dependent autoimmunity uniformly suppresses intestinal immune cell dissemination or instead changes which immune populations reach peripheral tissues. We therefore quantified the major lymphoid and myeloid populations among intestinal emigrants across the MLN, blood, spleen, and kidney (Fig. S2A). Lineage-specific changes were then analyzed across the MLN and blood (Table S1), spleen (Fig. 2A), and kidney (Fig. S2B-H). This analysis revealed distinct lineage- and tissue-dependent patterns. B and T cell dissemination generally decreased with increasing disease severity, whereas dendritic cell (DC) dissemination increased, particularly to the spleen and kidney, indicating that the overall reduction in gut-derived cells reflects selective reconfiguration rather than uniform loss of intestinal emigrants.

**Figure 2.**
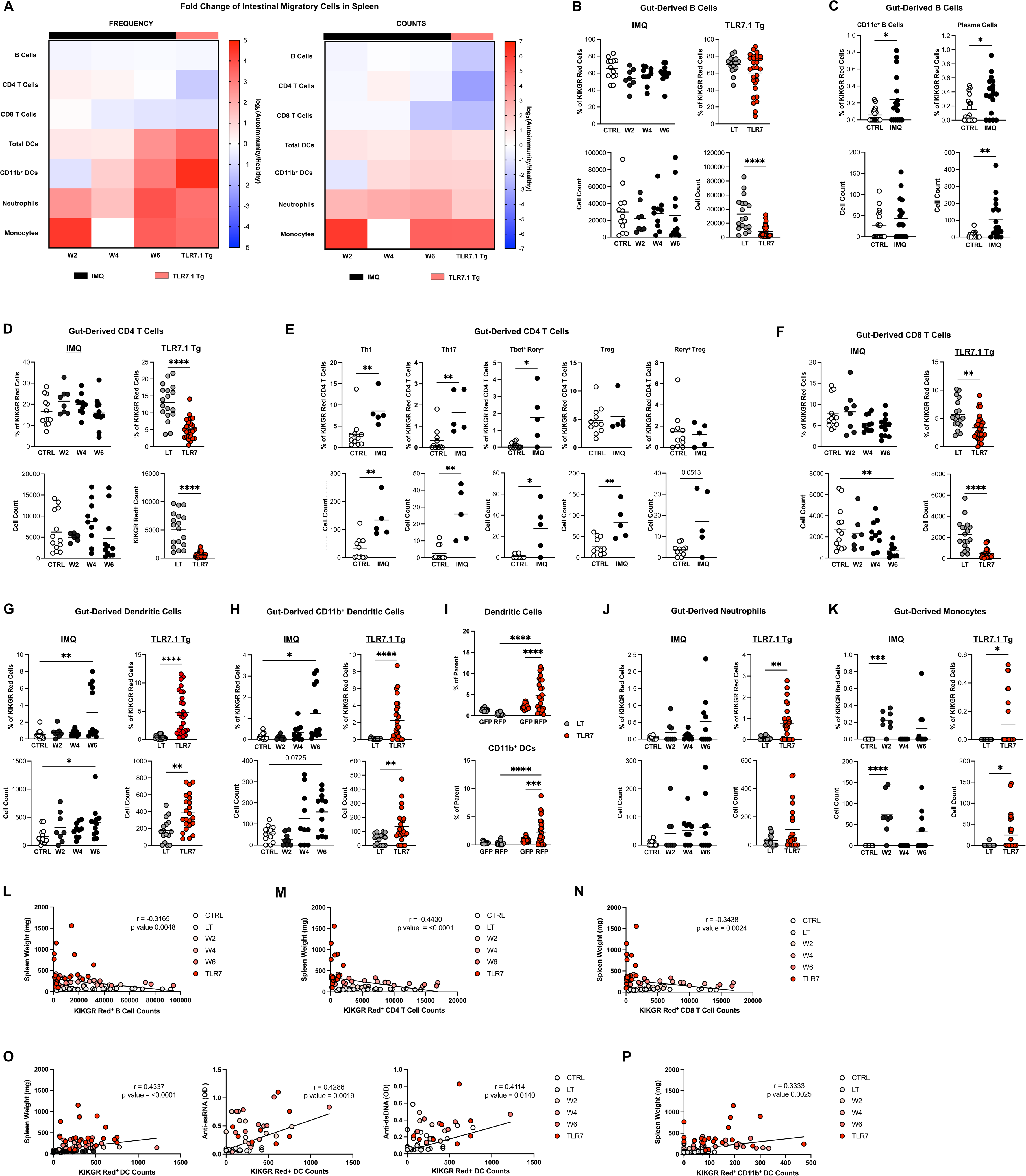
Systemic autoimmunity selectively reconfigures the cellular composition of intestinal emigrants. (A) Heatmaps showing fold change frequency and absolute counts of cell subsets of the intestinal migratory cells (RFP^+^) in the spleen. Values represent IMQ-treated KikGR mice at weeks 2, 4 and 6 (W2, W4, W6) relative to healthy KikGR controls (CTRL), and in KT adult mice (TLR7.1Tg) relative to their healthy littermate controls (LT). (B) Frequencies and absolute counts of B cells, (D) CD4 T cells, (F) CD8 T cells, (G) dendritic cells (DCs), (H) CD11b^+^ DCs, (J) neutrophils, and (K) monocytes out of the KikGR Red cells (RFP^+^) in IMQ-treated KikGR mice at weeks 2, 4 and 6 and in KT adult mice (TLR7.1Tg) in the Spleen (n = 8-29 per group). (C) Frequencies and absolute counts of CD11c^+^ B cells and Plasma Cells in IMQ-treated KikGR mice at week 6 of treatment (n = 17-18 per group). (E) Frequencies and absolute counts of CD4 T cell subsets Th1, Th17, T-bet⁺RORγt⁺, Tregs and RORγt⁺ Tregs out of KikGR Red CD4 T cells in IMQ-treated KikGR mice at week 6 of treatment (n = 5-11 per group). (I) Comparison of frequencies of dendritic cells and CD11b^+^ dendritic cells in KikGR Red (RFP^+^) and KikGR Green (GFP^+^) cells of KT adult mice (TLR7.1Tg) in the Spleen (n = 16-24 per group). (L) Correlation between gut-derived B cell absolute counts (KikGR Red^+^ B cells), (M) gut-derived CD4 T cell absolute counts (KikGR Red^+^ CD4 T cells), (N) gut-derived CD8 T cell absolute counts (KikGR Red^+^ CD8 T cells), and (P) CD11b^+^ dendritic cells (KikGR Red^+^ CD11b^+^ DCs) in the spleen and spleen weights, in IMQ-treated KikGR mice at weeks 2, 4 and 6 and in KT adult mice (TLR7.1Tg) (n = 8-23 per group). (O) Correlation between gut-derived dendritic cell absolute counts (KikGR Red^+^ dendritic cells) in the spleen and spleen weights, and anti-ssRNA and anti-dsDNA autoantibody OD values in IMQ-treated KikGR mice at weeks 2, 4 and 6 and in KT adult mice (TLR7.1Tg) (n = 8-23 per group). Data are shown as means. Statistical significance was determined by Kruskal-Wallis test with Dunn’s multiple comparison test (for more than two groups) and unpaired two-tailed Mann-Whitney U test (for two groups). Correlations were assessed using Spearman’s rank correlation. *\*p < 0.05, **p < 0.01, ***p < 0.001*.

B cells represented the predominant intestinal immune cell population disseminating to peripheral tissues under healthy conditions (Fig. 2A, B). However, their contribution to the migratory pool progressively decreased as TLR7-dependent autoimmunity became more severe. During IMQ-induced autoimmunity, B cell dissemination decreased in a tissue-dependent manner, with reduced B cell frequency in the MLN and blood at later stages and reduced dissemination to the kidney, whereas splenic dissemination was comparatively preserved (Fig. 2B; Fig. S2B; Table S1). In adult KT mice, gut-derived B cell numbers were markedly reduced in the spleen and kidney, with decreased frequencies particularly evident in the kidney (Fig. 2B; Fig. S2B; Table S1). Despite this loss, the phenotype of the B cells that continued to disseminate was substantially shifted. At 6 weeks of IMQ treatment, splenic gut-derived B cells showed increased frequencies of CD11c⁺ B cells ^25,26^ (Fig. 2C). Gut-derived plasma cells were also increased in both frequency and number in the spleen at 6 weeks of IMQ treatment (Fig. 2C). Thus, autoimmune progression reduced B cell dissemination but increased the representation of CD11c⁺ B cells among the remaining emigrants and, separately, increased gut-derived plasma-cell dissemination.

Gut-derived T cells showed a related but distinct pattern. Changes in total CD4⁺ T cell dissemination during IMQ treatment were comparatively modest and tissue-dependent, whereas adult KT mice exhibited a pronounced reduction in gut-derived CD4⁺ T cells, with fewer cells reaching the spleen and kidney and a marked decrease in their frequency in the spleen, while frequencies in the MLN and blood were comparatively preserved (Fig. 2D; Fig. S2C; Table S1). Importantly, phenotypic changes were already evident during IMQ-induced disease despite the more limited reduction in total CD4⁺ dissemination. By week 6, splenic gut-derived CD4⁺ T cells contained increased frequencies and numbers of Th1, Th17, and T-bet⁺RORγt⁺ cells, together with increased numbers of Treg cells (Fig. 2E). In contrast, RORγt⁺ Treg cells showed no significant increase. Gut-derived CD8⁺ T cell dissemination was also reduced with disease progression. By week 6 of IMQ treatment, CD8⁺ emigrant frequency was decreased in the MLN, while splenic dissemination was also reduced. In adult KT mice, CD8⁺ T cell frequency was reduced in the MLN, and substantially fewer gut-derived CD8⁺ T cells reached the spleen and kidney, while blood dissemination remained comparatively preserved (Fig. 2F; Fig. S2D; Table S1). These findings indicate that autoimmune progression progressively restricts bulk lymphocyte dissemination while selectively enriching the pool of intestinal migratory cells reaching the spleen for inflammatory CD4⁺ T cells.

In striking contrast to the progressive loss of lymphoid emigrants, intestinal dendritic cell dissemination increased during TLR7-dependent autoimmunity. Among the immune populations examined, DCs showed the most consistent increase in dissemination as disease progressed. Gut-derived dendritic cells progressively accumulated in the spleen and kidney during IMQ treatment, with increased frequency and number also evident in the spleens of adult KT mice (Fig. 2G; Fig. S2E). CD11b⁺ DCs showed a similar pattern, with increased dissemination to the spleen and kidney during IMQ-induced disease and prominent accumulation in the spleen of adult KT mice (Fig. 2H; Fig. S2F). In adult KT mice, both total DCs and CD11b⁺ DCs were markedly enriched within the gut-derived RFP⁺ compartment compared with non-gut-derived GFP⁺ cells, whereas no comparable enrichment was observed in healthy littermate controls (Fig. 2I). This preferential representation indicates that the increased splenic DC population during established autoimmunity is specifically associated with intestinal emigrants rather than simply reflecting generalized systemic DC expansion. Other myeloid populations also showed increased systemic dissemination, although these effects were more restricted. Monocyte dissemination increased transiently to the MLN and spleen during IMQ treatment, whereas in adult KT mice a significant increase was restricted to the spleen, with no change in the kidney (Fig. 2K; Fig. S2H; Table S1). Neutrophil dissemination showed only a trend toward increased splenic migration during IMQ treatment but was significantly increased in the spleen of adult KT mice, with no significant changes in the MLN, blood, or kidney (Fig. 2J; Fig. S2G; Table S1). Consistent with this lineage divergence, splenic dissemination of gut-derived B cells, CD4⁺ T cells, and CD8⁺ T cells correlated inversely with splenic enlargement across disease progression, whereas total DC and CD11b⁺ DC dissemination correlated positively with spleen weight. Total gut-derived DC abundance also correlated positively with anti-ssRNA and anti-dsDNA autoantibody levels (Fig. 2L-P).

Together, these findings reveal a marked divergence between the lymphoid and myeloid components of intestinal immune dissemination during systemic autoimmunity. Whereas dissemination of B and T lymphocytes progressively contracted with increasing disease severity, myeloid emigrants were maintained or increased, with dendritic cells showing the most consistent expansion in systemic tissues. The opposing relationships of lymphocyte and DC dissemination with disease severity further indicate that this shift occurs progressively as systemic pathology increases. Thus, TLR7-dependent autoimmunity does not simply suppress intestinal immune dissemination but progressively shifts the composition of the intestinal emigrant pool from predominantly lymphoid toward increased representation of myeloid cells. We next asked whether this divergent lymphoid-myeloid pattern was already present before the development of established systemic pathology.

### TLR7-dependent remodeling of intestinal immune dissemination precedes established systemic pathology

Because adult KT mice exhibited pronounced suppression and reorganization of intestinal immune dissemination, we next asked whether these changes emerged only after systemic pathology was established or were already present earlier during spontaneous TLR7-dependent autoimmunity. We therefore examined KT mice at 4 weeks of age, considerably before the adult stage analyzed above (Fig. 3A). At this age, KT mice showed no detectable splenic enlargement compared with age-matched littermate controls, in marked contrast to the pronounced splenomegaly observed in adult KT mice (Fig. S3A). Nevertheless, 4-week-old KT mice already exhibited significantly increased anti-dsDNA and anti-ssRNA autoantibodies (Fig. S3B). Thus, these mice enabled us to examine intestinal immune dissemination following the emergence of serological autoimmunity but prior to splenomegaly.

**Figure 3.**
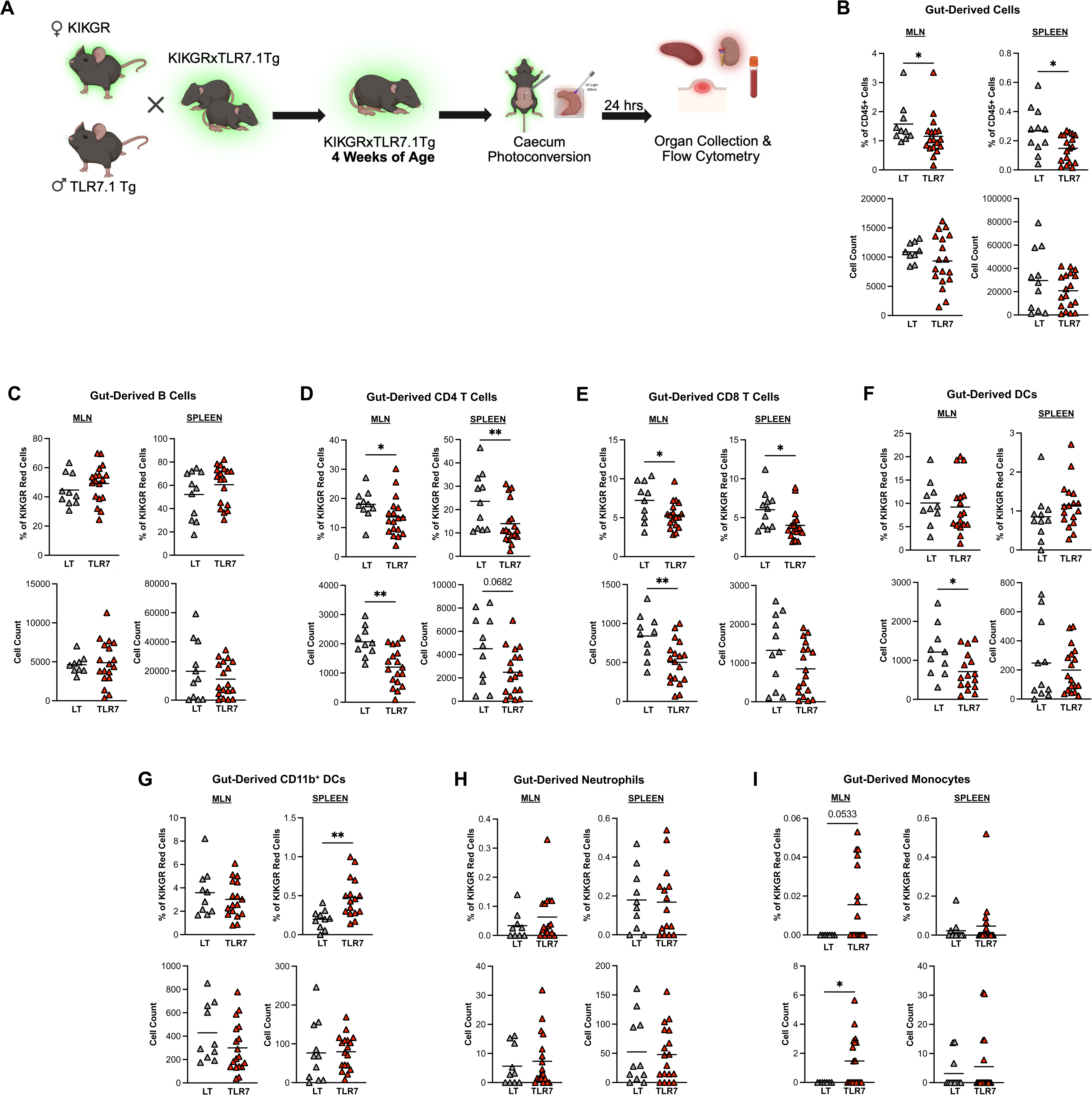
TLR7-dependent remodeling of intestinal immune dissemination precedes established systemic pathology. (A) Experimental design for photoconversion in young TLR7.1Tg model. For TLR7.1Tg, KikGR Tg mice were crossed with TLR7.1Tg to obtain KikGRxTLR7.1Tg; these mice underwent photoconversion at 4 weeks of age, and euthanasia and tissue collection were performed 24 hours after photoconversion. (B) Gut-derived RFP^+^ cell frequency and absolute counts in young KT mice (TLR7.1Tg) in the MLN and spleen. (C) Frequencies and absolute counts of B cells, (D) CD4 T cells, (E) CD8 T cells, (F) Dendritic cells (DCs), (G) CD11b^+^ DCs, (H) neutrophils, and (I) monocytes out of the KikGR Red cells (RFP^+^) young KT (TLR7.1Tg) at 4 weeks of age in the MLN and spleen (B-I, n = 9-18 per group). Data are shown as means. Statistical significance was determined by unpaired two-tailed Mann-Whitney U test. *\*p < 0.05, **p < 0.01, ***p < 0.001*.

Despite the absence of detectable splenomegaly, the frequency of gut-derived RFP⁺ cells was significantly decreased in both the MLN and spleen compared with age-matched littermate controls, although total RFP⁺ cell numbers were not significantly reduced in either tissue (Fig. 3B). In contrast, neither the frequency nor number of total gut-derived RFP⁺ cells differed significantly in the blood or kidney at this age (Fig. S3C), indicating that the early reduction in intestinal dissemination was initially concentrated within gut-associated and splenic compartments. Gut-derived B cell dissemination also remained largely preserved, with no significant differences in frequency or number in the MLN, blood, spleen, or kidney (Fig. 3C; Fig. S3D). Thus, the prominent reduction in B cell dissemination observed in adult KT mice had not yet been established by 4 weeks of age.

In contrast to the preserved B cell compartment, T cell dissemination was already substantially reduced. Gut-derived CD4⁺ T cells were decreased in frequency in both the MLN and spleen, accompanied by a significant reduction in CD4⁺ T cell numbers in the MLN and a similar trend in the spleen (Fig. 3D). In the blood, CD4⁺ T cell frequency and number were unchanged, whereas the kidney showed a trend toward reduced CD4⁺ T cell frequency without a significant change in cell number (Fig. S3E). Gut-derived CD8⁺ T cells showed a similar tissue-restricted pattern, with reduced frequency and number in the MLN and reduced frequency in the spleen (Fig. 3E), but no significant differences in the blood or kidney (Fig. S3F). Together, these findings indicate that reduced intestinal T cell dissemination is already evident in 4-week-old KT mice and is initially most pronounced in the MLN and spleen, before becoming more broadly distributed across systemic tissues in established disease.

In contrast to the broad reduction in T cell dissemination, myeloid populations did not show a comparable loss across tissues. Total gut-derived DC numbers were reduced in the MLN, but remained unchanged in the spleen, whereas the frequency of splenic CD11b⁺ DC emigrants was significantly increased in 4-week-old KT mice (Fig. 3F, G). DC and CD11b⁺ DC dissemination through the blood and kidney showed no significant increases at this age (Fig. S3G, H). Gut-derived monocyte numbers were significantly increased in the MLN, while no significant differences were detected in the blood, spleen, or kidney. Neutrophil dissemination similarly showed no significant changes across tissues, although upward trends were evident in selected blood and kidney measurements (Fig. 3H, I; Fig. S3I, J). Absolute numbers of gut-derived monocytes and neutrophils in these systemic compartments were very low, limiting interpretation of these trends. Thus, before splenomegaly, young KT mice already exhibited an early divergence between lymphoid and myeloid intestinal dissemination, characterized by reduced CD4⁺ and CD8⁺ T cell migration, along with selective increases in the frequencies of myeloid emigrants, most notably an increased representation of gut-derived CD11b⁺ DCs in the spleen. This pattern represents an early stage of the broader lymphoid-to-myeloid shift observed in adult disease and demonstrates that reorganization of intestinal immune dissemination precedes the establishment of systemic pathology, rather than arising solely from advanced autoimmune inflammation.

### The intestinal microbiota shapes physiological intestinal immune dissemination during health and autoimmunity

TLR7-dependent models of systemic autoimmunity are strongly influenced by changes in the intestinal microbiota, and lupus-associated pathobionts can worsen disease severity. We therefore asked whether the intestinal microbial environment also regulates the physiological dissemination of immune cells. We first compared untreated adult KikGR mice maintained in our standard colony with wild-type KikGR littermates generated within the KikGR x TLR7.1 breeding colony (Fig. 4A). Intestinal immune dissemination differed markedly between the two groups, with the strongest effects concentrated in the MLN (Fig. 4B-I). Wild-type mice from the TLR7.1 breeding colony showed a substantial increase in the number of total gut-derived RFP⁺ cells reaching the MLN. This increase included greater numbers of gut-derived B cells, CD4⁺ T cells, CD8⁺ T cells, and CD11b⁺ DCs, while their relative representation within the migratory pool was also reconfigured, with an increased B cell frequency and reduced CD4⁺ and CD8⁺ T cell frequencies (Fig. 4B-I). In contrast, dissemination to systemic tissues was not significantly altered, and the frequency of total and CD11b⁺ DCs in the kidney decreased (Table S2). Thus, chronic differences associated with the breeding environment strongly modified dissemination to the gut-draining MLN without producing a parallel increase in intestinal emigrants reaching systemic tissues.

**Figure 4.**
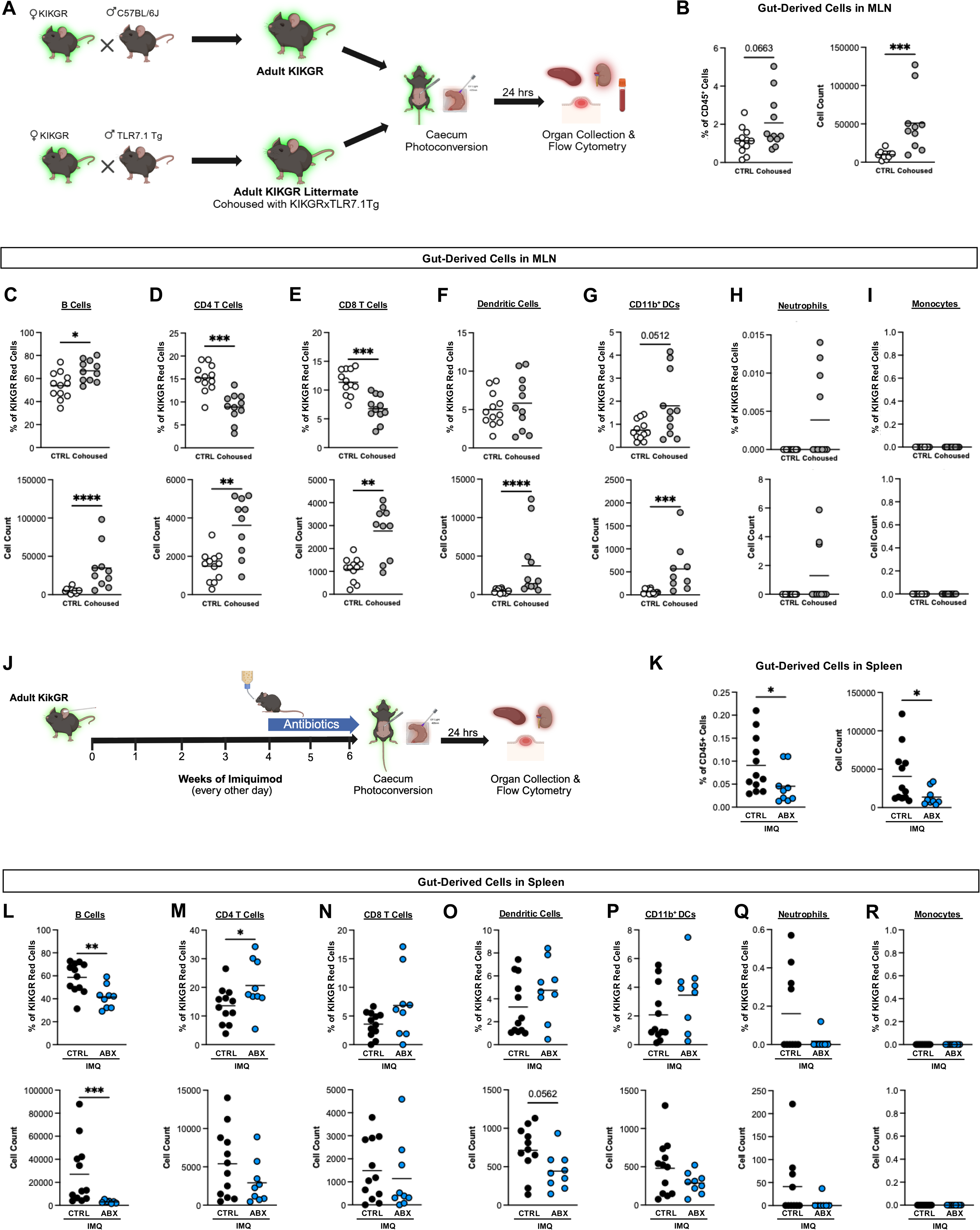
The intestinal microbiota shapes physiological intestinal immune dissemination during health and autoimmunity. (A) Experimental design for photoconversion in adult KikGR mice (CTRL) and adult KikGR littermate mice (Cohoused) cohoused with KT adults (KikGR x TLR7.1Tg). For the Cohoused group, we crossed KikGR mice with TLR7.1Tg to obtain KikGRxTLR7.1Tg and KikGR littermates (Cohoused). Mice underwent photoconversion at 12 weeks of age. In all cases, euthanasia and tissue collection were performed 24 hours after photoconversion. (B) Gut-derived RFP^+^ cell frequency and absolute counts in adult KikGR mice (CTRL) and adult KikGR littermate (Cohoused) in the MLN (n=11-12 per group). (C) Frequencies and absolute counts of B cells, (D) CD4 T cells, (E) CD8 T cells, (F) dendritic cells (DCs), (G) CD11b^+^ DCs, (H) neutrophils, and (I) monocytes out of the KikGR Red cells (RFP^+^) in adult KikGR mice (CTRL) and adult KikGR littermate (Cohoused) in the MLN (C-I; n=11-12 per group). (J) Experimental design for photoconversion in adult KikGR treated with and without antibiotics. All the mice were treated topically with IMQ three times a week for 6 weeks. Beginning at week 5, the control group (CTRL) received a single gavage of 200μl of ultra-pure water followed by autoclaved drinking water for 14 days, while the antibiotics group (ABX) received a single gavage of 200μl of streptomycin (20mg/mouse) followed by Ampicillin in drinking water (1g/L) for 14 days. Photoconversion was performed on the final day of water/antibiotic supplementation, and euthanasia and tissue collection were performed 24 hours after photoconversion. (K) Gut-derived RFP^+^ cell frequency and absolute counts in mice with (ABX) and without (CTRL) antibiotic supplementation in the spleen (n=9-12 per group). (L) Frequencies and absolute counts of B cells, (M) CD4 T cells, (N) CD8 T cells, (O) dendritic cells (DCs), (P) CD11b^+^ DCs, (Q) neutrophils, and (R) monocytes out of the KikGR Red cells (RFP^+^) in mice with (ABX) and without (CTRL) antibiotic supplementation in the spleen (L-R; n=9-12 per group). Data are shown as means. Statistical significance was determined by unpaired two-tailed Mann-Whitney U test. *\*p < 0.05, **p < 0.01, ***p < 0.001*.

We next asked whether short-term manipulation of the microbiota was sufficient to reproduce these effects in otherwise healthy KikGR mice. Fecal microbiota from either healthy wild-type or adult TLR7.1 Tg donors was transferred to KikGR recipients, which were analyzed two weeks later (Fig. S4A). Neither donor microbiota significantly changed total intestinal immune dissemination to the MLN, spleen, or kidney (Fig. S4B), and no broad changes were detected among the major gut-derived lymphoid and myeloid populations examined (Table S3). We also tested *Enterococcus gallinarum*, a lupus-associated intestinal pathobiont that exacerbates TLR7-dependent autoimmunity, using the related commensal *Enterococcus faecalis* as a control (Fig. S4C). Following repeated colonization over one week, total gut-derived RFP⁺ cell dissemination to the MLN, spleen, and kidney remained comparable among control, *E. faecalis*-, and *E. gallinarum*-colonized mice (Fig. S4D). Most migratory immune populations were similarly unchanged; however, *E. gallinarum* selectively increased the frequency of gut-derived DCs reaching the kidney at day 7 (Table S4). Thus, acute fecal microbiota transfer or short-term colonization with a lupus-associated pathobiont did not broadly modify systemic intestinal immune dissemination in the absence of ongoing autoimmune inflammation.

We previously showed that microbiota depletion reduced TLR7-dependent autoimmune pathology in two settings: IMQ-induced disease in germ-free mice, in which the microbiota was absent before and throughout disease induction, and TLR7.1 Tg mice following broad-spectrum antibiotic treatment initiated before disease onset ^16^. Here, we instead asked how antibiotic treatment affects intestinal immune dissemination when initiated during ongoing IMQ-induced autoimmunity. KikGR mice were treated with IMQ for 6 weeks and received streptomycin and ampicillin only during the final 2 weeks (weeks 4 to 6; Fig. 4J). Late antibiotic treatment significantly reduced splenomegaly but did not alter anti-ssRNA or anti-dsDNA autoantibody levels (Fig. S4E, F). Compared with IMQ treatment alone, antibiotic treatment significantly reduced both the frequency and number of total gut-derived RFP⁺ cells reaching the spleen (Fig. 4K). This reduction was driven prominently by B cells, whose frequency and absolute number among splenic intestinal emigrants were markedly decreased following antibiotic treatment (Fig. 4L). In contrast, the frequency of gut-derived CD4⁺ T cells in the spleen increased without a corresponding increase in cell number (Fig. 4M), while CD8⁺ T cell representation showed a similar upward trend in the MLN but remained unchanged in the spleen (Fig 4N; Fig. S4I). Splenic gut-derived DC numbers also showed a downward trend while other myeloid cells remained unchanged, whereas the frequency of CD11b⁺ DCs among kidney emigrants increased (Fig. 4O-4R; Fig. S4K). Additional lineage analyses in the MLN and kidney revealed more limited changes (Fig. S4G-M). Thus, antibiotic treatment during ongoing autoimmunity reduced splenomegaly without affecting autoantibody production, while further suppressing dissemination of selected intestinal populations, particularly splenic B cells, and shifting the cellular composition of the remaining migratory pool.

Together, these findings demonstrate that microbial regulation of intestinal immune dissemination depends strongly on inflammatory context and timing. Chronic environmental differences primarily affected dissemination within the gut-associated MLN, whereas acute microbiota transfer or pathobiont colonization produced limited systemic effects. In contrast, antibiotic treatment during ongoing TLR7-dependent autoimmunity substantially reconfigured intestinal immune dissemination, most prominently in the spleen. Thus, the influence of the intestinal microbiota on physiological immune dissemination becomes broader in the context of ongoing systemic autoimmune inflammation.

## DISCUSSION

Intestinal immune cell dissemination has emerged as a fundamental process that contributes to systemic immune homeostasis and can influence inflammatory disease through individual migratory immune populations ^2–6^. Here, we show that TLR7-dependent autoimmunity progressively suppresses this physiological dissemination pathway while selectively reconfiguring the cellular composition of intestinal emigrants. As disease severity increases, dissemination shifts from a predominantly lymphoid migratory pool toward greater representation of myeloid cells, with dendritic cells showing the most consistent increase in systemic tissues. Importantly, this lineage divergence is already evident in young mice predisposed to develop systemic autoimmunity, before detectable splenomegaly, indicating that reorganization of intestinal immune dissemination is not simply a consequence of advanced systemic inflammation. Together, these findings establish remodeling of physiological intestinal immune dissemination as an early and dynamic feature of TLR7-dependent autoimmunity.

The opposing behaviors of lymphoid and myeloid emigrants suggest that systemic autoimmunity does not uniformly increase or suppress intestinal trafficking but instead differentially reshapes the migratory programs of individual immune lineages. B cells were the dominant migratory population under healthy conditions but progressively declined with increasing disease severity, and the dissemination of CD4⁺ and CD8⁺ T cells was also reduced. However, this contraction of bulk lymphocyte dissemination was accompanied by selective enrichment of activated and effector populations. Splenic gut-derived CD11c⁺ B cells increased in frequency, gut-derived plasma cells increased in both frequency and number, and splenic gut-derived CD4⁺ T cells showed increased frequencies and numbers of Th1, Th17, and T-bet⁺RORγt⁺ effector populations during IMQ-induced disease. These findings suggest that autoimmunity restricts bulk lymphocyte dissemination without uniformly restricting activated or effector lymphoid populations. In contrast, DCs showed the most consistent increase in systemic dissemination, particularly to the spleen and kidney, with more restricted increases also observed among monocytes and neutrophils. Several non-mutually exclusive mechanisms could contribute to this divergence. Chronic intestinal antigen exposure or inflammation may favor lymphocyte retention, tissue residency, or altered responsiveness to migratory cues, while simultaneously promoting DC recruitment, activation, or export from the intestine. Consistent with the possibility that these lineage-specific changes arise within a broadly altered trafficking environment, ileal transcriptional profiling revealed extensive remodeling of chemokine ligands and receptors in both inducible and spontaneous TLR7-dependent autoimmunity. Which of these pathways drive the lineage-specific changes in dissemination observed here remains to be determined. The preferential representation of gut-derived DCs relative to non-gut-derived DCs in adult KT spleens further suggests that their accumulation is not merely a reflection of generalized systemic DC expansion. Notably, the early reduction in T cell dissemination together with increased splenic representation of intestinal CD11b⁺ DCs was already evident in young KT mice, before the broad dissemination defects observed in adults and before splenomegaly developed. These findings therefore support a model in which TLR7-dependent autoimmunity progressively reorganizes intestinal dissemination through lineage-specific migratory programs rather than through a uniform change in immune cell movement.

Our findings also reveal that microbial regulation of intestinal immune dissemination is strongly dependent on timing and inflammatory context. Even in healthy mice, intestinal immune dissemination was sensitive to microbial and environmental context. Chronic differences associated with the breeding environment and housing markedly altered dissemination to the gut-draining MLN, whereas short-term fecal microbiota transfer produced little broad-based change, and acute *E. gallinarum* colonization had comparatively limited effects aside from selective increases in DC dissemination. In contrast, antibiotic treatment initiated during ongoing IMQ-induced autoimmunity substantially reconfigured intestinal emigrant populations, most prominently in the spleen, and further reduced gut-derived B cell dissemination. Notably, even this relatively short period of antibiotic treatment reduced splenomegaly without altering anti-ssRNA or anti-dsDNA autoantibody levels, indicating that antibiotic-mediated perturbation of the microbiota can differentially affect components of established autoimmune pathology. This dissociation may reflect differences in the kinetics and microbial dependence of splenic inflammatory expansion and established humoral responses; splenic inflammatory expansion may respond rapidly to microbiota depletion, whereas pre-existing autoantibodies and antibody-producing populations may persist over the two-week intervention. Interestingly, young KT mice showed the converse temporal relationship, with autoantibodies emerging before spleen enlargement, further emphasizing that these manifestations of TLR7-dependent autoimmunity are not obligatorily coupled. This finding is consistent with our previous observations that the absence or earlier depletion of the microbiota attenuates TLR7-dependent disease ^16^, and shows that microbiota-targeting intervention can modify selected features of disease even after autoimmunity is established. Whether the accompanying changes in intestinal dissemination result directly from loss of microbial signals, from the selective reduction in systemic pathology, or from both remains unresolved. Nevertheless, these findings indicate that the microbiota and the host’s inflammatory state interact to determine how intestinal immune populations disseminate beyond the gut.

The functional consequences of this remodeling remain an important question. The preferential increase in intestinal DC dissemination suggests that chronic autoimmunity alters not only the number of cells leaving the gut but also the quantity and nature of intestinal information delivered to distant tissues. Gut-derived DCs could transport microbial antigens or products, host antigens, or inflammatory signals to the spleen, kidney, and potentially other immune sites, including the thymus. In early life, we previously showed that intestinal DCs can transport microbiota-derived antigens to the thymus and shape the development of microbiota-specific T cells ^4^, raising the possibility that systemic inflammation re-engages aspects of developmental gut-to-systemic immune communication later in life. Whether the enhanced DC dissemination observed here alters systemic or thymic T cell responses, promotes disease, represents an adaptive response, or simply accompanies inflammation will require functional testing. Similarly, the mechanisms underlying the early divergence between lymphocyte and DC dissemination remain unknown and will require defining the chemokine pathways, tissue-derived signals, and cell-intrinsic programs that control each migratory lineage. Understanding how these pathways are reorganized before the establishment of pathology, and whether they can be selectively manipulated, may reveal new strategies to modulate tissue-to-tissue immune communication during systemic autoimmunity.

### Limitations of the study

This study defines how physiological intestinal immune dissemination is remodeled during TLR7-dependent autoimmunity but does not establish the molecular mechanisms driving these lineage-specific migratory changes. Although we identify reduced lymphoid dissemination together with relative preservation or increased representation of myeloid emigrants, particularly dendritic cells, the chemokine pathways, tissue-derived signals, and cell-intrinsic programs responsible for these divergent responses remain to be determined. Likewise, while our findings identify intestinal dendritic cells and other migratory populations as candidates for influencing systemic immune responses, we did not directly assess the biological cargo they transport or determine whether their dissemination contributes causally to autoimmune pathogenesis. Finally, our microbiota perturbation experiments demonstrate that microbial regulation of intestinal immune dissemination depends on timing and inflammatory context, but do not identify the specific microbial factors, products, or metabolites responsible for these broader context-dependent effects.

## RESOURCE AVAILABILITY

### Lead contact

Further information and requests for resources and reagents should be directed to the Lead Contact, Dr. Daniel Zegarra-Ruiz, who will fulfill them.

### Materials availability

This study did not generate new unique reagents.

### Data and code availability

The mRNA-seq data generated in this study were deposited in the NCBI Gene Expression Omnibus (GEO) database under accession GSE315533. This paper does not report any original code. Any data requests will be fulfilled by the lead contact.

## ACKNOWLEDGMENTS

Flow cytometry and cell sorting were performed in the University of Virginia (UVA) Flow Cytometry Core Facility (RRID: SCR_017829), supported in part by NCI grant P30-CA044579; cell sorting on the Sony MA900 CS was enabled through NIH S10 instrumentation grant 1S10OD034355-01. This work was supported by a National Institute of Arthritis and Musculoskeletal and Skin Diseases R00 grant (R00AR080207), the UVA TransUniversity Microbiome Initiative (TUMI) Quick Turnaround Grant, and the Empowering Lupus Research Career Development Award from the Lupus Research Alliance. We thank Jessica Hamerman (Benaroya Research Institute) for TLR7.1 transgenic mice and Martin Kriegel (University of Münster) for *Enterococcus gallinarum* isolates. We are also grateful to Gretchen Diehl and Amanda Chen (Memorial Sloan Kettering Cancer Center) for guidance on the use of KikGR mice. Finally, we thank members of our laboratory, as well as members of the Beirne B. Carter Center for Immunology Research and the Department of Microbiology, Immunology, and Cancer Biology at UVA, for helpful feedback throughout this study.

## AUTHOR CONTRIBUTIONS

Conceptualization and supervision were performed by D.F.Z.-R., who also led funding acquisition, visualization, and writing of the original draft and revisions. Data acquisition was carried out by K.S.S.-M., S.R.D.-P., M.K., and O.B. Data analysis was conducted by K.S.S.-M., M.K., and M.V.-A. Visualization and figure preparation were performed by K.S.S.-M., M.V.-A., and D.F.Z.-R. The original manuscript was written by D.F.Z.-R., and reviewed and edited by K.S.S.-M., S.R.D.-P., M.K., and D.F.Z.-R.

## DECLARATION OF GENERATIVE AI AND AI-ASSISTED TECHNOLOGIES

During the preparation of this work, the authors used ChatGPT (OpenAI) for editing, proofreading, and language optimization. The authors reviewed and edited all content as needed and take full responsibility for the content of the publication

## SUPPLEMENTARY FIGURES AND TABLES

**Figure S1.** Experimental validation and complementary analyses of TLR7-dependent intestinal immune dissemination

**Figure S2.** Extended characterization of intestinal emigrants during TLR7-dependent autoimmunity

**Figure S3.** Complementary analyses of intestinal immune dissemination in young KT mice

**Figure S4.** Complementary analyses of microbiota-dependent intestinal immune dissemination

**Table S1.** Extended analysis of intestinal emigrant populations in the MLN and blood during TLR7-dependent autoimmunity

**Table S2.** Complementary analysis of intestinal immune dissemination associated with the breeding environment

**Table S3.** Intestinal immune dissemination following fecal microbiota transfer

**Table S4.** Intestinal immune dissemination following short-term pathobiont colonization

## STAR★METHODS

### EXPERIMENTAL MODEL AND STUDY PARTICIPANT DETAILS

#### Mice

C57BL/6J (B6, JAX 000664), Tg(CAG-KikGR)33Hadj/J (KikGR, JAX 013753) ^24^, TLR7.1 transgenic (TLR7.1 Tg) lupus-prone mice were used in this study. TLR7.1 Tg mice were kindly provided by Jessica Hamerman (Benaroya Research Institute) ^17^. All other strains were originally obtained from The Jackson Laboratory and subsequently bred in-house under specific pathogen-free (SPF) conditions at the University of Virginia vivarium. All mouse strains were maintained on a C57BL/6J genetic background. Mice were used between 3 and 15 weeks of age unless otherwise indicated. Experiments included both male and female mice, and age- and sex-matched animals were used whenever possible. Littermate controls were used when available, and animals were randomly assigned to experimental groups. Mice were provided standard laboratory chow and water ad libitum and housed on hardwood chip bedding in a temperature-controlled, air-conditioned facility under a 12-hour light/dark cycle. All procedures were conducted in accordance with protocols approved by the University of Virginia Institutional Animal Care and Use Committee.

### METHOD DETAILS

#### Lupus models

##### Inducible lupus model

Mice received a topical treatment of 25mg of 5% Imiquimod (IMQ) cream (Perrigo) on the ears, alternating between right and left, three times a week for 6 weeks, which induces a lupus-like disease as previously described. This protocol is adapted from Zegarra-Ruiz et al., 2019 ^16,18^.

##### Spontaneous lupus model

TLR7.1 Tg mice were used as a spontaneous model of lupus-like disease. These mice exhibit a genetic predisposition to autoimmunity, characterized by approximately 8–16-fold increases in TLR7 expression, leading to spontaneous onset of systemic lupus erythematosus-like manifestations, including splenomegaly, autoantibody production, and inflammatory pathology such as glomerulonephritis, beginning around 6 weeks of age (Zegarra et al., 2019) ^16,17^. Mice were analyzed at 4 weeks of age and again at 12 weeks of age.

#### Photoconversion and intestinal immune cell tracking (KikGR Mice)

The cecum of Tg(CAG-KikGR)33Hadj/J (KikGR, JAX 013753) mice was photoconverted using a 405 nm violet laser (peak power, 5 mW) applied for 5 minutes. Mice were anesthetized with 2.5% isoflurane (vol/vol) in 2 L/min oxygen and maintained at 37°C for the duration of the procedure. A 2 cm midline abdominal incision was performed to exteriorize the cecum, which was continuously hydrated with sterile saline. Following photoconversion, the cecum was repositioned into the abdominal cavity, and the incision was closed with sutures. Twenty-four hours after photoconversion, tissues were harvested and analyzed by flow cytometry to detect photoconverted KIKGR Red⁺ cells.

#### Bacterial Stock Preparation and Storage

*Enterococcus gallinarum ^12^* was cultured anaerobically at 37°C in pre-reduced Brain Heart Infusion (BHI) (Millipore Sigma) medium and expanded to the exponential phase (∼2 × 10^9^ CFU). Cultures were pooled, mixed 1:1 with BHI (60%) supplemented with glycerol (40%) and L-cysteine (0.1%) (Millipore Sigma), aliquoted, and cryopreserved at −80°C.

*Enterococcus faecalis ATCC 29212* was cultured anaerobically at 37°C in pre-reduced Brain Heart Infusion (BHI) (Millipore Sigma) medium and expanded to the exponential phase (∼1 × 10^9^ CFU). Cultures were pooled, mixed 1:1 with BHI (60%) supplemented with glycerol (40%) (Millipore Sigma), aliquoted, and cryopreserved at −80°C.

#### Bacterial Colonization

Mice were colonized with either *Enterococcus gallinarum or Enterococcus faecalis* using frozen stocks (∼1 × 10^9^ CFU). Colonization was performed 3 times a week by oral gavage (2 × 10^8^ CFU per mouse, per gavage).

*E. gallinarum and E. faecalis* colonization were confirmed by culture from fecal pellets. Fecal material was homogenized through a 100 µm strainer, resuspended in sterile PBS, and inoculated into BHI medium supplemented with vancomycin (6 µg/mL) (Millipore Sigma), including pure culture and uninoculated controls. Cultures were incubated anaerobically at 37°C for 72 hours, pelleted, washed twice with sterile PBS, and resuspended prior to DNA isolation. For DNA isolation, bacterial cells were pelleted, washed in TE buffer (10 mM Tris-HCl pH 8.0, 1 mM EDTA), and resuspended. Cells were treated with lysozyme (2.5 mg/mL) (Millipore Sigma), 20% SDS, and proteinase K (0.9 mg/mL) (Millipore Sigma), followed by phenol–chloroform extraction. DNA was precipitated with ethanol and sodium acetate, washed, air-dried, and resuspended in nuclease-free water (Thermo Fisher Scientific).

Bacterial abundance was assessed by conventional PCR using species-specific primers:

*E. gallinarum* (F: TTACTTGCTGATTTTGATTCG; R: TGAATTCTTCTTTGAAATCAG) ^12^

*E. faecalis* (F: CCGAGTGCTTGCACTCAATTGG; R: CTCTTATGCCATGCGGCATAAAC)

#### Antibiotics administration

At the beginning of week 5 of imiquimod treatment, mice received a single dose of Streptomycin (200mg/mL, 100 μL per mouse) by oral gavage. On the same day, ampicillin (1g/L) was added to the drinking water for a 14-day period ^27^ (week 5 and week 6 of imiquimod treatment).

#### Fecal Microbiota Transplant (FMT)

Fecal pellets were collected from adult TLR7.1 Tg mice, homogenized, and filtered in 1mL of PBS through a 70-μm strainer (3 fecal pellets per 5 recipient mice). Recipient Tg(CAG-KikGR)33Hadj/J received 200 μL of the resulting suspension by oral gavage. Control mice received a transplant prepared from their own fecal pellets following the same method. FMT was performed 3 times a week for 2 weeks.

#### Tissue processing and cell isolation

##### Cell isolation

Single-cell suspensions were prepared from blood, spleen, kidney, mesenteric lymph node (MLN) and large intestine lamina propria.

Blood was collected in EDTA-coated tubes before euthanasia and centrifuged at 1,800 rpm for 5 min at 4°C. Plasma was discarded and red blood cells were lysed using RBC lysis buffer (BioLegend) for 5 minutes at room temperature, and centrifuge at 1,800 rpm for 5 min at 4°C, this process was repeated 3 times before resuspending in MACS buffer (PBS containing 2% FBS (Gibco) and 2 mM EDTA (Thermo Fisher Scientific)).

Following euthanasia, the abdominal cavity was opened, and tissues were carefully collected. Tissues other than the large intestine were mechanically dissociated in digestion buffer (complete RPMI supplemented with 0.01% v/v of 150 mg/mL DNase I from bovine pancreas (Millipore Sigma) and 0.01% v/v of 100 kU/mL Type 8 Collagenase from *Clostridium histolyticum* (Millipore Sigma)) and digested for 30 min at 37°C. After digestion, cell suspensions were filtered through 70-µm strainers, washed in 10ml of MACS buffer, centrifuged at 1,800 rpm for 5 min at 4°C, and resuspended in MACS buffer unless otherwise indicated. For spleen and kidney, red blood cells were lysed using RBC lysis buffer (BioLegend) for 5 minutes at room temperature. Kidney cell suspensions were subsequently enriched by 40%/80% Percoll density gradient centrifugation (Cytiva) at 2,500 rpm for 20 min at room temperature without brake. Cells from the interphase were collected, washed, and resuspended in MACS buffer. Cell suspensions were filtered through 70-µm strainers and processed as described above.

To isolate cells from the large intestinal lamina propria, intestines were harvested and placed in ice-cold PBS. Adipose tissue removed, intestines were opened longitudinally, and luminal contents were discarded. Tissues were cut into approximately 2-cm segments and washed in PBS. Epithelial cells and mucus were removed by two sequential incubations at 37°C with shaking (200 rpm): first in PBS containing 1 mM dithiothreitol (DTT) (Thermo Fisher Scientific), 30 mM ethylenediaminetetraacetic acid (EDTA) (Thermo Fisher Scientific), and 10 mM HEPES for 10 min, followed by a second incubation in PBS containing 30 mM EDTA and 10 mM HEPES (Fisher Scientific) for 10 min. Tissues were then digested in Type 8 collagenase/DNase I-containing digestion buffer in complete RPMI for 1 h at 37°C with shaking. Following digestion, cells were collected and enriched by 40%/80% Percoll density gradient centrifugation as described for kidney samples. Cells were washed and resuspended in MACS buffer for downstream applications.

#### Flow cytometry

Cells isolated from spleen, kidneys, MLN, small intestinal lamina propria, large intestine lamina propria, or blood were surface-stained in MACS buffer or Fc/Block buffer for 25 minutes in the dark at 4 °C with fluorescently conjugated antibodies.

For the majority of experiments, cells were stained with a comprehensive extracellular antibody panel designed to identify major immune populations. We used antibodies specific to CD4 (1:300, BioLegend, cat #: 100516), CD8a (1:300, BioLegend, cat #: 100752), TCRβ (1:200, BioLegend, cat #: 109241), CD11b (1:400, BioLegend, cat #: 101226), CD11c (1:400, BioLegend, cat #: 117328), Ly6G (1:400, BioLegend, cat #: 127648), I-A/I-E (1:400, BioLegend, cat #: 107641), B220 (1:300, BioLegend, cat #: 103222), CD45 (1:400, BioLegend, cat #: 103149), Ly6C (1:200, BioLegend, cat #: 128024) and PDCA-1 (1:300, BioLegend, cat #: 127039), Viability was assessed using DAPI (1:300, BioLegend, cat #:422801).

Data were acquired the same day using an Attune NxT flow cytometer (Thermo Fisher Scientific) and analyzed using FlowJo v10 (Tree Star). Absolute cell numbers were determined using Precision Count Beads (BioLegend).

##### T cell phenotyping

For T cells, we used antibodies specific to CD3 (1:200, BioLegend, cat #: 100229), CD4 (1:300, BioLegend, cat #: 100548), CD8a (1:300, BioLegend, cat #: 100752), TCRβ (1:200, BioLegend, cat #: 109241), and CD45 (1:300, BioLegend, cat #: 103149).

For identification of T cell subsets, Zombie Violet Fixable Viability Kit (BioLegend cat #: 423114) was included in the extracellular antibody cocktail. Then, cells were stained according to the published dish soap protocol to maintain the RFP and GFP signal ^28^. For intracellular staining, cells were stained with antibodies against FOXP3 (1:100, eBioscience, cat #: 56-5773-82), RORγt (1:100, eBioscience, cat #: 25-6981-82), T-bet (1:100, BioLegend, cat #: 644828),) overnight at 4 °C.

Data were acquired using an Attune NxT flow cytometer (Thermo Fisher Scientific) and analyzed using FlowJo v10 (Tree Star). Absolute cell numbers were determined using Precision Count Beads (BioLegend).

##### B cell phenotyping

For B Cell phenotyping, antibodies against CD11c (1:400, BioLegend, cat #: 117328), I-A/I-E (1:400, BioLegend, cat #: 107641), CD19 (1:300, BioLegend, cat #: 115521), B220 (1:300, BioLegend, cat #: 103244), CD45 (1:300, BioLegend, cat #: 103149), CX3CR1 (1:200, BioLegend, cat #: 149027), CD90.2 (1:300, BioLegend, cat #: 140318), F4/80 (1:300, BioLegend, cat #: 123133) and CD138 (1:200, BioLegend, cat #: 142521). Viability was assessed using DAPI (1:300, BioLegend, cat #: 422801).

Data were acquired the same day using an Attune NxT flow cytometer (Thermo Fisher Scientific) and analyzed using FlowJo v10 (Tree Star). Absolute cell numbers were determined using Precision Count Beads (BioLegend).

##### Gating strategy: T cells

After exclusion of dead cells and doublets, KIKGR Red^+^ cells were defined as CD45^+^ RFP^+^, from KIKGR Red^+^, CD8 T cells were defined as CD3^+^ TCRβ^+^ CD8^+^ and CD4 T cells as CD3^+^ TCRβ^+^ CD4^+^. Th1, Th17, regulatory T cell (Treg), and RORγt^+^ Treg populations were identified as T-bet^+^ (Th1), RORγt^+^ (Th17), FOXP3^+^ (Tregs), RORγt^+^ FOXP3^+^ (RORγt^+^ Tregs) and Tbet^+^ RORγt^+^ (Tbet^+^ RORγt^+^)

##### Gating strategy: B cells

After exclusion of dead cells and doublets, KIKGR Red^+^ cells were defined as CD45^+^ RFP^+^, from KIKGR Red^+^, B cells were identified as CD90.2^-^ F4/80^-^ CX3CR1^-^ CD19⁺ B220⁺ cells, plasma cells were defined as CD90.2^-^ F4/80^-^ CX3CR1^-^ CD138^+^ cells. Autoimmune B cells were defined as CD90.2^-^ F4/80^-^ CX3CR1^-^ CD19⁺ B220⁺ CD11c^+^.

##### Gating strategy: Multilineage panel

After exclusion of dead cells and doublets, KIKGR Red^+^ cells were defined as CD45^+^ RFP^+^, from KIKGR Red^+^, DCs were defined as CD45^+^ RFP^+^CD11c⁺ MHCII⁺ cells and subdivided into CD11b^+^ and CD11b^-^ subsets. Plasmacytoid Dendritic Cells (pDCs) were defined as CD45^+^RFP^+^TCR^-^B220^+^CD11c^int^PDCA-1^+^. Monocytes were defined as CD45^+^RFP^+^TCR^-^B220^-^CD11c^-^MHCII⁻CD11b⁺Ly6C+ and Neutrophils as CD45^+^RFP^+^TCR^-^B220^-^CD11c^-^MHCII⁻CD11b⁺Ly6G^+^.

##### RNA extraction and RNA sequencing

Ileum sections were collected in TRIzol (Thermo Fisher Scientific) containing 1.0 mm zirconium-silicate beads (BioSpec Products) and either stored at −80°C or processed immediately. Samples were lysed by bead beating (3 × 60 s, maximum speed) with cooling on ice between cycles. RNA was extracted by chloroform (Millipore Sigma) phase separation, and the aqueous phase was precipitated with isopropanol (Millipore Sigma), washed twice with 75% ethanol, briefly air-dried, and resuspended in ultrapure water, according to the manufacturer’s protocol. RNA concentration and purity were assessed using a NanoDrop. RNA libraries were prepared using poly(A) selection and sequenced by Genewiz (Azenta Life Sciences) on an Illumina NovaSeq platform in 2×150 bp paired-end mode, yielding approximately 20 million read pairs per sample.

##### RNA-seq preprocessing and exploratory analysis

Raw paired-end reads were quality-trimmed using fastp (v0.23.4). Adapter sequences were autodetected and removed, polyG tails were trimmed, and low-quality bases were filtered using a sliding window approach (window size 4, mean quality threshold 24). Reads with a Phred quality score below 15 or shorter than 50 bp after trimming were discarded. Read quality before and after trimming was assessed using FastQC and aggregated with MultiQC. Transcript quantification was performed from cleaned reads using Salmon (v1.10.3) in mapping-based mode with automatic library type detection. The reference index was built from the GENCODE mouse transcriptome (release vM36, GRCm39 assembly) using the Salmon index. Transcript-level abundance estimates were aggregated to gene-level counts using the tximport R package with gene-to-transcript mappings derived from the GENCODE vM36 annotation file.

##### Differential gene expression analysis and functional enrichment analyses

Differential expression analysis was performed using DESeq2 (v1.46.0). Samples were modeled based on experimental condition (CTRL, IMQ, TLR7), and pairwise comparisons were extracted from a single joint model: IMQ vs. CTRL, TLR7 vs. CTRL, and TLR7 vs. IMQ. Size factor normalization and dispersion estimation were performed using standard DESeq2 settings. Differential expression results were extracted using Wald tests with Benjamini-Hochberg correction for multiple hypothesis testing. Genes were considered significantly differentially expressed based on an adjusted p-value (padj) <0.1 and an absolute log2fold change >1.

Heatmaps of variance-stabilized transformed (VST) expression values were generated using ComplexHeatmap. Gene-wise z-scores were calculated for row-scaled visualization of relative expression across samples. Rows were hierarchically clustered, while columns were split by condition.

#### Enzyme-linked immunosorbent assays (ELISAs)

Autoantibodies against double-stranded DNA (dsDNA) and single-stranded RNA (ssRNA) in serum were quantified by ELISA assay.

##### Anti-dsDNA ELISA

Plates (High Binding, Corning) were pre-coated with protamine sulfate (100 µL per well at 150 µg/mL) and incubated for 45 minutes at room temperature in the dark. Wells were rinsed twice with ultrapure water, blotted, and air-dried. Plates were then coated with 100 µg/mL salmon sperm DNA (Thermo Fisher Scientific) and blocked with PBS containing 3% BSA and 5% fetal bovine serum (FBS). Serum samples were diluted 1:1000 in dilution buffer and incubated for 1 hour at 37°C. Wells were washed and incubated with rabbit anti–mouse IgG conjugated to HRP (0.1 µg/mL, Fisher Scientific), for 1 hour at 37°C. Color development was performed using TMB substrate, and the reaction was stopped with 2M sulfuric acid.

##### Anti-ssRNA ELISA

Plates were pre-coated with protamine sulfate (100 µL per well at 50 µg/mL) and processed as above. Plates were coated with 5 mg/mL yeast RNA (Thermo Fisher Scientific) and blocked with PBS containing 0.05% Tween-20 and 5% FBS. Serum samples were diluted 1:1000 in dilution buffer and incubated for 1 hour at 37°C. Subsequent washing, secondary antibody incubation, and development steps were performed as described for the anti-dsDNA ELISA.

Absorbance was measured at 450 nm with a 570 nm reference wavelength.

### QUANTIFICATION AND STATISTICAL ANALYSIS

All statistical analyses were performed using GraphPad Prism (v10.6.1). Individual data points are shown alongside the mean. “n” represents the number of biological samples analyzed in each experiment and the value of n is indicated in the corresponding figure legend. Differences between two groups were evaluated using a two-tailed unpaired Mann-Whitney U test. For comparisons involving more than two groups, a Kruskal-Wallis test followed by Dunn’s multiple-comparison test was used. Correlations were assessed using Spearman’s rank correlation. Outliers were identified using the ROUT method (Q = 1%) and excluded prior to statistical analysis. A P value < 0.05 was considered statistically significant. Statistical significance is indicated as follows: ns, not significant; *P ≤ 0.05; **P ≤ 0.01; ***P ≤ 0.001; and ****P ≤ 0.0001.

## REFERENCES

1. Morton, A.M., Sefik, E., Upadhyay, R., Weissleder, R., Benoist, C., and Mathis, D. (2014). Endoscopic photoconversion reveals unexpectedly broad leukocyte trafficking to and from the gut. Proc. Natl. Acad. Sci. U. S. A. 111, 6696–6701. 10.1073/pnas.1405634111.

2. Galván-Peña, S., Zhu, Y., Hanna, B.S., Mathis, D., and Benoist, C. (2024). A dynamic atlas of immunocyte migration from the gut. Sci Immunol 9, eadi0672. 10.1126/sciimmunol.adi0672.

3. Ennamorati, M., Vasudevan, C., Clerkin, K., Halvorsen, S., Verma, S., Ibrahim, S., Prosper, S., Porter, C., Yeliseyev, V., Kim, M., et al. (2020). Intestinal microbes influence development of thymic lymphocytes in early life. Proc. Natl. Acad. Sci. U. S. A. 117, 2570–2578. 10.1073/pnas.1915047117.

4. Zegarra-Ruiz, D.F., Kim, D. V., Norwood, K., Kim, M., Wu, W.J.H., Saldana-Morales, F.B., Hill, A.A., Majumdar, S., Orozco, S., Bell, R., et al. (2021). Thymic development of gut-microbiota-specific T cells. Nature 594, 413–417. 10.1038/s41586-021-03531-1.

5. Fan, T., Tai, C., Sleiman, K.C., Cutcliffe, M.P., Kim, H., Liu, Y., Li, J., Xin, G., Grashel, M., Baert, L., et al. (2025). Aberrant T follicular helper cells generated by TH17 cell plasticity in the gut promote extraintestinal autoimmunity. Nat. Immunol. 26, 790–804. 10.1038/s41590-025-02125-7.

6. Teng, F., Klinger, C.N., Felix, K.M., Bradley, C.P., Wu, E., Tran, N.L., Umesaki, Y., and Wu, H.J.J. (2016). Gut Microbiota Drive Autoimmune Arthritis by Promoting Differentiation and Migration of Peyer’s Patch T Follicular Helper Cells. Immunity 44, 875–888. 10.1016/j.immuni.2016.03.013.

7. Pöysti, S., Toivonen, R., Takeda, A., Silojärvi, S., Yatkin, E., Miyasaka, M., and Hänninen, A. (2022). Infection with the enteric pathogen C. rodentium promotes islet-specific autoimmunity by activating a lymphatic route from the gut to pancreatic lymph node. Mucosal Immunol. 15, 471–479. 10.1038/s41385-022-00490-2.

8. Brea, D., Poon, C., Benakis, C., Lubitz, G., Murphy, M., Iadecola, C., and Anrather, J. (2021). Stroke affects intestinal immune cell trafficking to the central nervous system. Brain Behav. Immun. 96, 295–302. 10.1016/j.bbi.2021.05.008.

9. Ghafouri-Fard, S., Shahir, M., Taheri, M., and Salimi, A. (2021). A review on the role of chemokines in the pathogenesis of systemic lupus erythematosus. Cytokine 146, 155640. 10.1016/j.cyto.2021.155640.

10. Dziedzic, R., Siwiec-Koźlik, A., Koźlik-Siwiec, P., Węglarczyk, K., Siedlar, M., Korkosz, M., and Kosałka-Węgiel, J. (2026). Serum Levels of Selected Chemokines in Patients with Systemic Lupus Erythematosus Correlate with Disease Activity and Clinical Features: Results from a Single-Center Observational Study. Biomedicines 14. 10.3390/biomedicines14030567.

11. Tsokos, G.C. (2020). Autoimmunity and organ damage in systemic lupus erythematosus. Nat. Immunol. 21, 605–614. 10.1038/s41590-020-0677-6.

12. Manfredo Vieira, S., Hiltensperger, M., Kumar, V., Zegarra-Ruiz, D., Dehner, C., Khan, N., Costa, F.R.C., Tiniakou, E., Greiling, T., Ruff, W., et al. (2018). Translocation of a gut pathobiont drives autoimmunity in mice and humans. Science (1979). 359, 1156–1161. 10.1126/science.aar7201.

13. Fine, R.L., Manfredo Vieira, S., Gilmore, M.S., and Kriegel, M.A. (2020). Mechanisms and consequences of gut commensal translocation in chronic diseases. Gut Microbes 11, 217–230. 10.1080/19490976.2019.1629236.

14. Azzouz, D., Omarbekova, A., Heguy, A., Schwudke, D., Gisch, N., Rovin, B.H., Caricchio, R., Buyon, J.P., Alekseyenko, A. V., and Silverman, G.J. (2019). Lupus nephritis is linked to disease-activity associated expansions and immunity to a gut commensal. Ann. Rheum. Dis. 78, 947–956. 10.1136/annrheumdis-2018-214856.

15. Kume, M., Din, J., and Zegarra-Ruiz, D.F. (2025). Dysregulated Intestinal Host–Microbe Interactions in Systemic Lupus Erythematosus: Insights from Patients and Mouse Models. Microorganisms 13. 10.3390/microorganisms13030556.

16. Zegarra-Ruiz, D.F., El Beidaq, A., Iñiguez, A.J., Lubrano Di Ricco, M., Manfredo Vieira, S., Ruff, W.E., Mubiru, D., Fine, R.L., Sterpka, J., Greiling, T.M., et al. (2019). A Diet-Sensitive Commensal Lactobacillus Strain Mediates TLR7-Dependent Systemic Autoimmunity. Cell Host Microbe 25, 113–127.e6. 10.1016/j.chom.2018.11.009.

17. Deane, J.A., Pisitkun, P., Barrett, R.S., Feigenbaum, L., Town, T., Ward, J.M.M., Flavell, R.A., and Bolland, S. (2007). Control of Toll-like Receptor 7 Expression Is Essential to Restrict Autoimmunity and Dendritic Cell Proliferation. Immunity 27, 801–810. 10.1016/j.immuni.2007.09.009.

18. Yokogawa, M., Takaishi, M., Nakajima, K., Kamijima, R., Fujimoto, C., Kataoka, S., Terada, Y., and Sano, S. (2014). Epicutaneous application of toll-like receptor 7 agonists leads to systemic autoimmunity in wild-type mice: A new model of systemic lupus erythematosus. Arthritis and Rheumatology 66, 694–706. 10.1002/art.38298.

19. Brown, G.J., Cañete, P.F., Wang, H., Medhavy, A., Bones, J., Roco, J.A., He, Y., Qin, Y., Cappello, J., Ellyard, J.I., et al. (2022). TLR7 gain-of-function genetic variation causes human lupus. Nature 605, 349–356. 10.1038/s41586-022-04642-z.

20. Sun, X., Chao, J.L., Gerner, M., and Elkon, K.B. (2025). Induction of Type I Interferon–Dependent Activation and Migration of Inflammatory Dendritic Cells to Local Lymph Nodes by UV Light Exposure. Arthritis and Rheumatology 77, 867–875. 10.1002/art.43108.

21. Skopelja-Gardner, S., Tai, J., Sun, X., Tanaka, L., Kuchenbecker, J.A., Snyder, J.M., Kubes, P., Mustelin, T., Elkon, K.B., and Snyder, J. (2021). Acute skin exposure to ultraviolet light triggers neutrophil-mediated kidney inflammation. Proc Natl Acad Sci U S A. 118, e2019097118. 10.1073/pnas.2019097118.

22. Liao, X., Pirapakaran, T., and Luo, X.M. (2016). Chemokines and Chemokine Receptors in the Development of Lupus Nephritis. Mediators Inflamm. 2016, 6012715. 10.1155/2016/6012715.

23. Goel, R.R., Wang, X., O, L.J., Nakabo, S., Hasneen, K., Gupta, S., Wigerblad, G., Blanco, L.P., Kopp, J.B., Morasso, M.I., et al. Interferon lambda promotes immune dysregulation and tissue inflammation in TLR7-induced lupus. 10.1073/pnas.1916897117/-/DCSupplemental.

24. Nowotschin, S., and Hadjantonakis, A.K. (2009). Use of KikGR a photoconvertible green-to-red fluorescent protein for cell labeling and lineage analysis in ES cells and mouse embryos. BMC Dev. Biol. 9. 10.1186/1471-213X-9-49.

25. Golinski, M.-L., Demeules, M., Derambure, C., Riou, G., Maho-Vaillant, M., Boyer, O., Joly, P., and Calbo, S. (2020). CD11c+ B Cells Are Mainly Memory Cells, Precursors of Antibody Secreting Cells in Healthy Donors. Front. Immunol. 11. 10.3389/fimmu.2020.00032.

26. Wang, S., Wang, J., Kumar, V., Karnell, J.L., Naiman, B., Gross, P.S., Rahman, S., Zerrouki, K., Hanna, R., Morehouse, C., et al. (2018). IL-21 drives expansion and plasma cell differentiation of autoreactive CD11chiT-bet+ B cells in SLE. Nat. Commun. 9, 1758. 10.1038/s41467-018-03750-7.

27. Kim, M., Galan, C., Hill, A.A., Wu, W.J., Fehlner-Peach, H., Song, H.W., Schady, D., Bettini, M.L., Simpson, K.W., Longman, R.S., et al. (2018). Critical Role for the Microbiota in CX 3 CR1 + Intestinal Mononuclear Phagocyte Regulation of Intestinal T Cell Responses. Immunity 49, 151–163.e5. 10.1016/j.immuni.2018.05.009.

28. Burton, O.T., Dooley, J., and Liston, A. (2025). Introducing the Dish Soap Protocol: A Unified Approach for Multi-Modal Intracellular Staining. Curr. Protoc. 5. 10.1002/cpz1.70206.

