## Supplementary material for "TLR7-Dependent Autoimmunity Remodels Intestinal Immune Cell Dissemination": Supp_figures

### Figure S1. Experimental validation and complementary analyses of TLR7-dependent intestinal immune dissemination

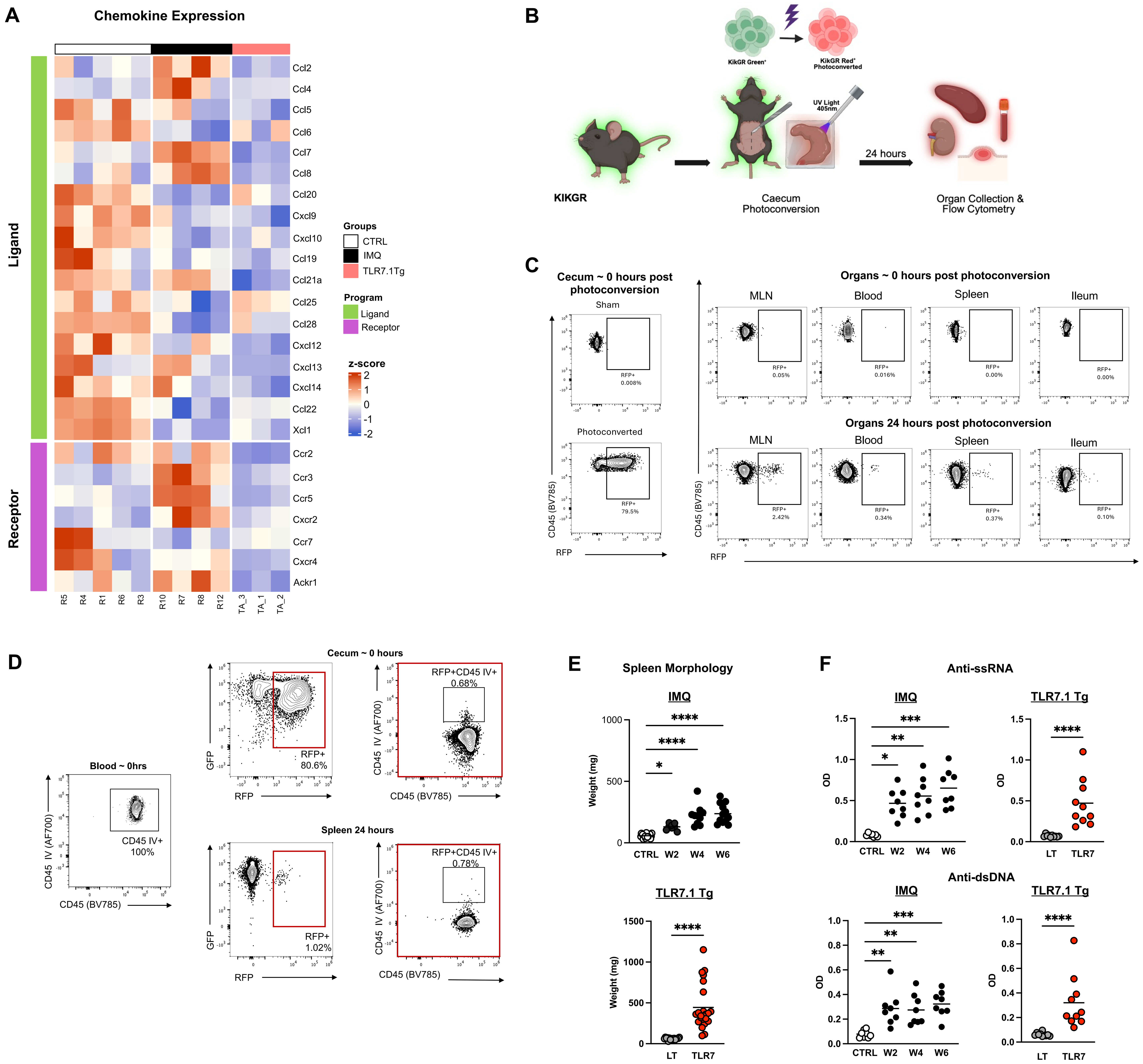

**Figure S1: Experimental validation and complementary analyses of TLR7-dependent intestinal immune dissemination**

(A) Heatmap showing chemokine (ligands and receptors) expression changes in B6 control mice (CTRL), mice treated with imiquimod for 6 weeks (IMQ), and adult TLR7.1Tg mice (TLR7.1Tg) (n = 3-5 per group). (B) Experimental design of photoconversion in KikGR mice. (C) Flow cytometry representative plots of cecum KikGR Red (RFP+) cells in non-photoconverted and photoconverted tissue immediately after the surgery, and flow cytometry representative plots of MLN, blood, spleen, and ileum KikGR Red (RFP+) cells immediately and 24 hours after photoconversion. (D) Flow cytometry representative plots of intravenously injected CD45 in cecum immediately after photoconversion and in the spleen 24 hours after photoconversion. (E) Spleen weights (n=8-29 per group) and (F) autoantibodies anti-ssRNA and anti-dsDNA in serum (n=8-10 per group) in IMQ-treated KikGR mice at weeks 2, 4 and 6 and in KT adult mice (TLR7.1Tg).

**Figure S2. Extended characterization of intestinal emigrants during TLR7-dependent autoimmunity**

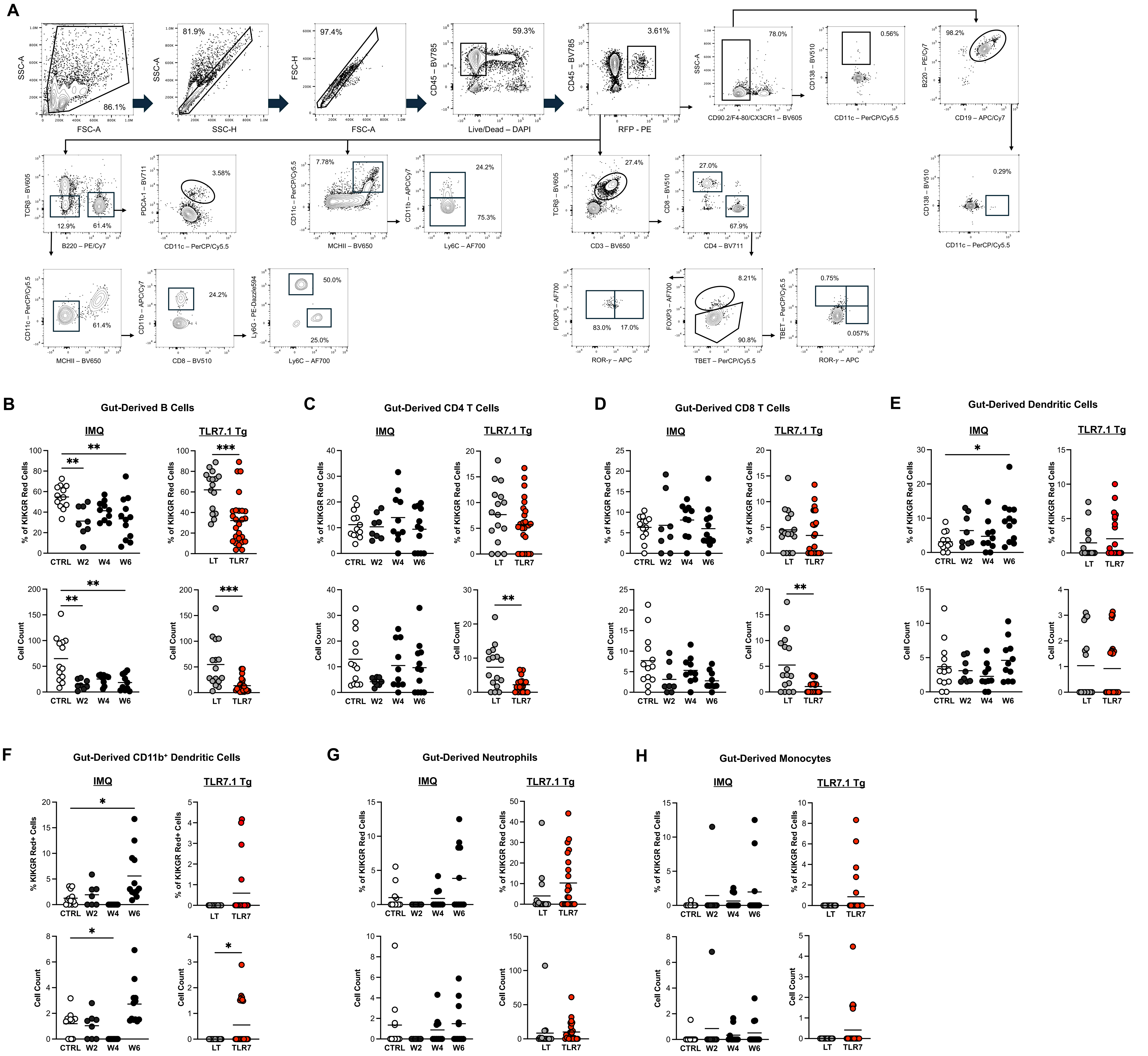

**Figure S2: Extended characterization of intestinal emigrants during TLR7-dependent autoimmunity**

(A) Gating strategy for KikGR Red cells (RFP+) and their downstream immune subsets. Following exclusion of debris and doublets (FSC-A/SSC-A, SSC-A/SSC-H, FSC-H/FSC-A), live cells were gated as Live/Dead (DAPI)- within CD45-BV785+ events, and photoconverted cells were identified as RFP-PE+. In the B cell branch, a dump gate (CD90.2/F4-80/CX3CR1-BV605-) was used to exclude T cells, macrophages, and monocytes, followed by CD138-BV510+ (plasma cells) and CD19-APC/Cy7+ vs B220-PE/Cy7+ to identify B cells, CD11c-PerCP/cy5.5+ B cells were also evaluated. In parallel, cells were resolved by TCRβ-BV605 vs B220-PE/Cy7 to separate T cells from other populations: non-T/B cells were further defined by PDCA-1 vs CD11c-PerCP/Cy5.5 (pDCs), CD11c vs MHCII-BV650+ (DCs), followed by CD11b-APC/Cy7 (CD11b$^{+}$ DCs); and CD11c vs MHCII-BV650- followed by Ly6C-AF700/ Ly6G-PE-Dazzle594 to distinguish monocytes and neutrophils. T cells (TCRβ+CD3+) were gated by CD4-BV711 vs CD8-BV510, and CD4 T cells were further resolved into Treg/Th17/Th1 subsets using FOXP3-AF700 vs TBET-PerCP/Cy5.5. (B) Frequencies and absolute counts of B cells, (C) CD4 T cells, (D) CD8 T cells, (E) dendritic cells (DCs), (F) CD11b$^{+}$ DCs, (G) neutrophils, and (H) monocytes out of the KikGR Red cells (RFP+) in IMQ-treated KikGR mice at weeks 2, 4 and 6 and in KT adult mice (TLR7.1Tg) in the kidney (n = 8-29 per group).

### Figure S3: Complementary analyses of intestinal immune dissemination in young KT mice

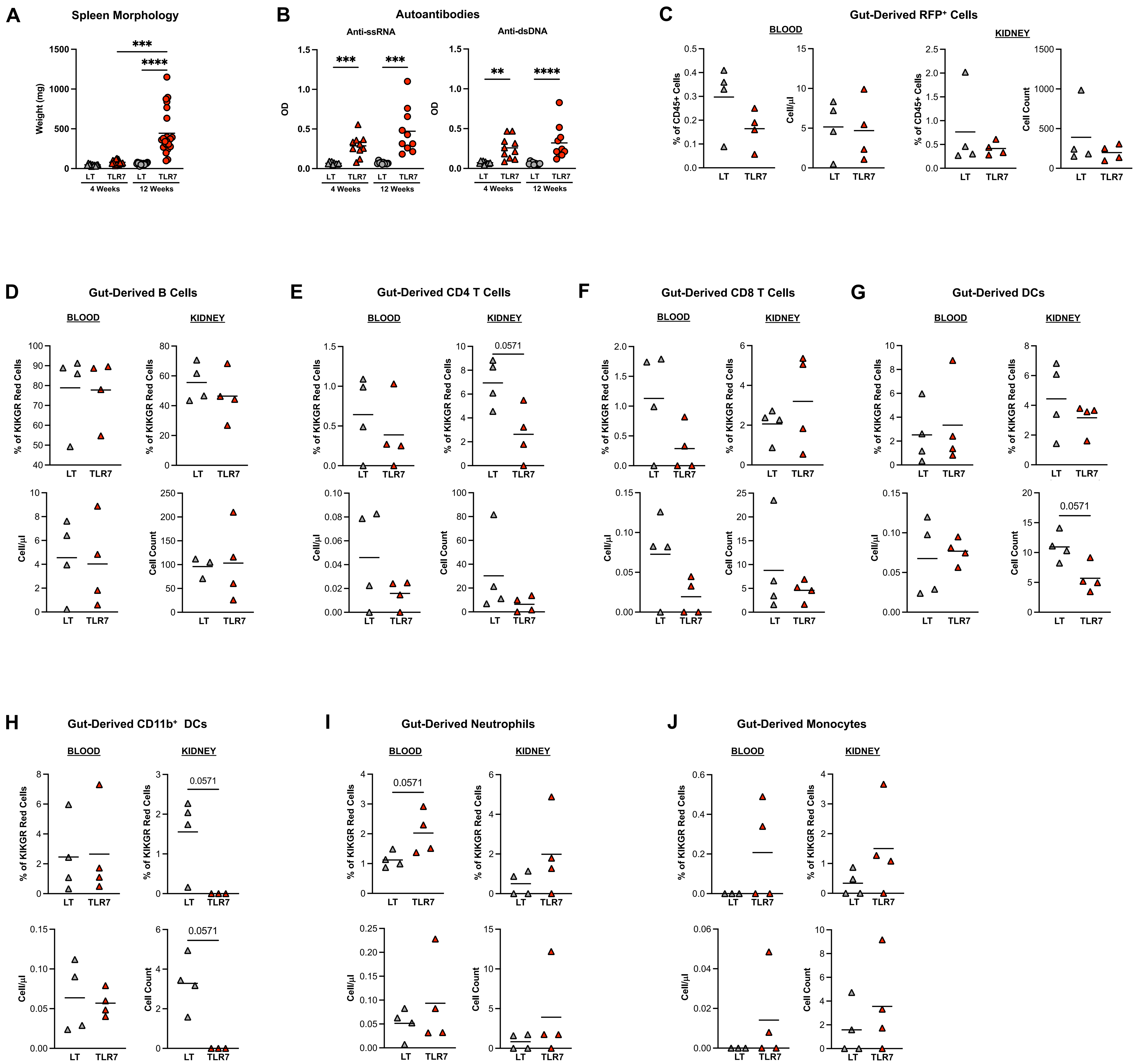

**Figure S3: Complementary analyses of intestinal immune dissemination in young KT mice**

(A) Spleen weight and (B) autoantibodies anti-ssRNA and anti-dsDNA in serum in young KT mice (TLR7.1Tg at 4 weeks of age) and adult KT mice (TLR7.1Tg at 12 weeks of age) compared to age-matched KikGR littermates. (C) Gut-derived RFP+ cell frequency and absolute counts in young KT mice (TLR7.1Tg) in blood and kidney. (D) Frequencies and absolute counts of B cells, (E) CD4 T cells, (F) CD8 T cells, (G) Dendritic cells (DCs), (H) CD11b$^{+}$ DCs, (I) neutrophils, and (J) monocytes out of the KikGR Red cells (RFP+) young KT (TLR7.1Tg) at 4 weeks of age in blood and kidney (C-J, n = 3-4 per group).

### Figure S4. Complementary analyses of microbiota-dependent intestinal immune dissemination

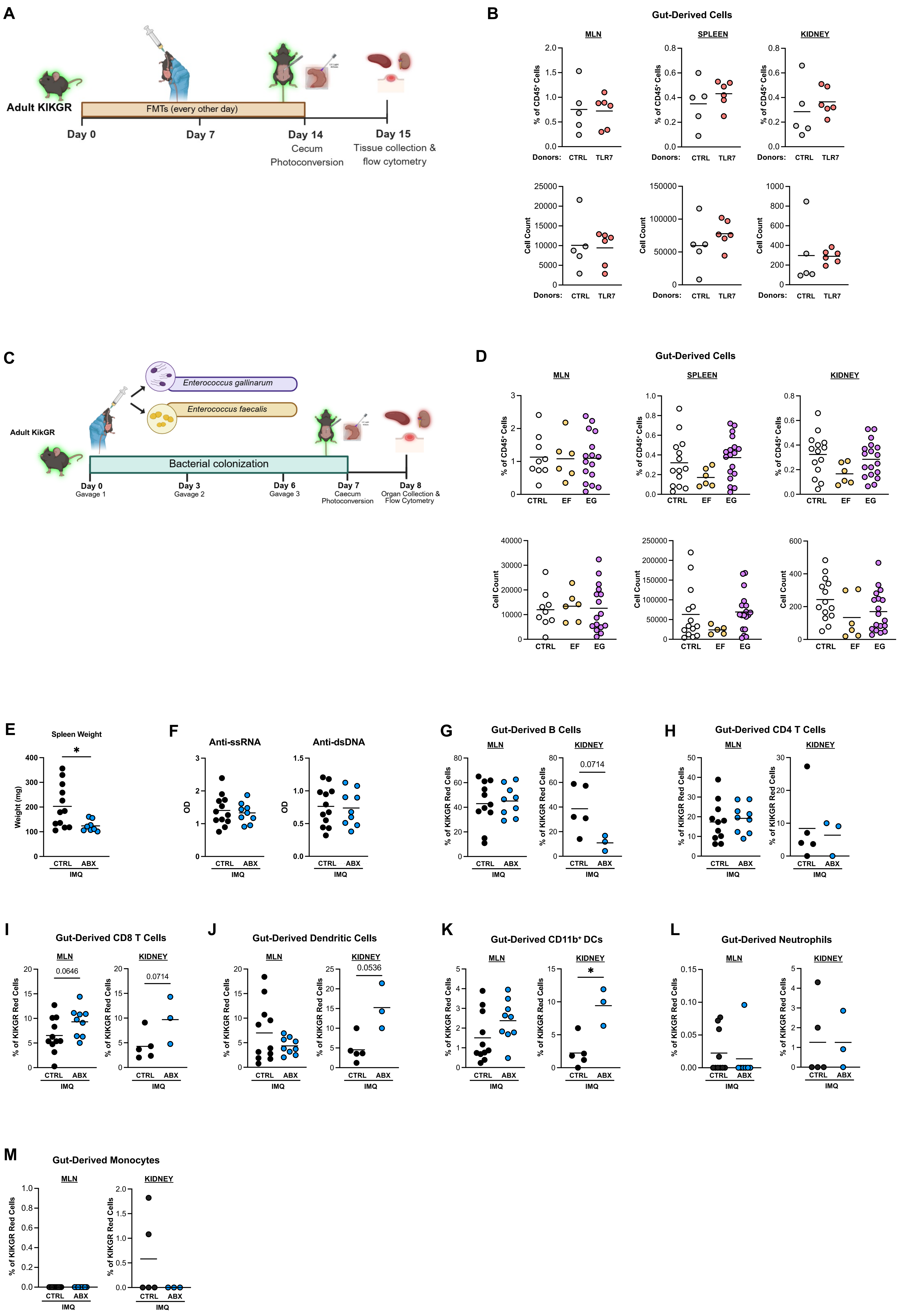

**Figure S4. Complementary analyses of microbiota-dependent intestinal immune dissemination**

(A) Experimental design for photoconversion in adult KikGR mice treated with fecal microbiota transplant (FMT) every other day for 14 days. Recipients KikGR received 200 µL FMT suspension by oral gavage , prepared either from their own fecal pellets (FMT CTRL) or from adult TLR7.1Tg donors (FMT TLR7). Photoconversion was performed on day 14 and euthanasia and tissue collection were performed 24 hours after photoconversion. (B) Gut-derived RFP+ cell frequency and absolute counts in KikGR mice treated with fecal microbiota transplant (FMT) in MLN, spleen, and kidney (n=5-6 per group). (C) Experimental design for photoconversion in adult KikGR mice colonized either with *Enterococcus gallinarum* (EG) or *Enterococcus faecalis* (EF) three times for a week by oral gavage. The control group (CTRL) was gavaged with PBS three times for a week. Photoconversion was performed on day 7, and euthanasia and tissue collection were performed 24 hours after photoconversion. (D) Gut-derived RFP+ cell frequency and absolute counts in KikGR mice colonized either with *Enterococcus gallinarum* (EG) or *Enterococcus faecalis* (EF) in MLN, spleen, and kidney (n=6-19 per group). (E) Spleen weight and (F) autoantibodies anti-ssRNA and anti-dsDNA in serum in mice with (ABX) and without (CTRL) antibiotic supplementation (n = 8-12 per group). (G) Frequency of B cells, (H) CD4 T cells, (I) CD8 T cells, (J) Dendritic cells (DCs), (K) CD11b$^{+}$ DCs, (L) neutrophils, and (M) monocytes in mice with (ABX) and without (CTRL) antibiotic supplementation in MLN and kidney (G-M, n = 3-12 per group).
