## Supplementary material for "TLR7-Dependent Autoimmunity Remodels Intestinal Immune Cell Dissemination": Table1

**Table S1. Extended analysis of intestinal emigrant populations in the MLN and blood during TLR7-dependent autoimmunity**

| **B Cells** | **Control** | **W2** | **W4** | **W6** | **LT** | **TLR7.1Tg** |
|---|---|---|---|---|---|---|
| Blood | 86.910±3.695 | 67.680±7.873 | 83.590±2.294 | 56.950±8.647[a] | 89.700±2.359 | 82.680±3.884 |
| MLN | 53.790±3.313 | 39.300±6.393 | 35.940±4.597[a] | 57.560±4.106 | 66.700±2.773 | 59.860±3.884 |
| **CD4 T Cells** | **Control** | **W2** | **W4** | **W6** | **LT** | **TLR7.1Tg** |
| Blood | 0.984±0.391 | 1.900±0.565 | 3.077±0.959 | 3.964±1.525 | 0.326±0.116 | 0.114±0.083 |
| MLN | 15.220±0.858 | 19.310±2.180 | 20.240±1.740 | 17.110±2.130 | 8.877±0.926 | 8.285±0.707 |
| **CD8 T Cells** | **Control** | **W2** | **W4** | **W6** | **LT** | **TLR7.1Tg** |
| Blood | 0.890±0.295 | 1.724±0.777 | 2.191±0.707 | 1.505±0.515 | 0.729±0.245 | 0.738±0.298 |
| MLN | 11.360±0.642 | 11.360±1.814 | 11.270±1.356 | 6.973±1.384[a] | 6.764±0.689 | 4.254±0.476[b] |
| **Monocytes** | **Control** | **W2** | **W4** | **W6** | **LT** | **TLR7.1Tg** |
| Blood | 0.000±0.000 | 0.188±0.119 | 0.000±0.000 | 0.169±0.813[a] | 0.000±0.000 | 0.000±0.000 |
| MLN | 0.000±0.000 | 0.064±0.025[a] | 0.087±0.040[a] | 0.022±0.012 | 0.002±0.001 | 0.025±0.009 |
| **Neutrophils** | **Control** | **W2** | **W4** | **W6** | **LT** | **TLR7.1Tg** |
| Blood | 0.442±0.226 | 2.714±1.477 | 0.598±0.234 | 1.646±0.630 | 0.728±0.291 | 3.489±1.476 |
| MLN | 0.000±0.000 | 0.013±0.007[a] | 0.000±0.000 | 0.000±0.000 | 0.004±0.002 | 0.018±0.007 |
| **Dendritic Cells ($MHCII^+$ $CD11c^+$)** | **Control** | **W2** | **W4** | **W6** | **LT** | **TLR7.1Tg** |
| Blood | 0.552±0.165 | 2.769±1.026 | 0.971±0.252 | 1.214±0.366 | 0.596±0.231 | 1.022±0.392 |
| MLN | 4.993±0.634 | 9.096±2.822 | 8.851±1.817 | 1.259±0.371[a] | 5.828±1.040 | 7.595±1.092 |
| **$CD11b^+$ DCs** | **Control** | **W2** | **W4** | **W6** | **LT** | **TLR7.1Tg** |
| Blood | 0.303±0.111 | 1.527±0.936 | 0.463±0.157 | 1.881±0.869 | 0.101±0.048 | 0.000±0.000 |
| MLN | 0.743±0.121 | 3.014±0.746[a] | 2.321±0.459 | 0.330±0.073 | 1.803±0.420 | 2.563±0.388 |

Values represent frequencies (% of KikGR $Red^+$ cells) and are expressed as mean±SEM (n=7-12 mice per group).
[a]= significantly different from control group (Kruskal-Wallis test followed by Dunn’s multiple-comparison test and a P<0.05 was considered statistically significant).
[b]= significantly different from Littermate (LT) group.(Mann-Whitney test was performed for this comparison and a P<0.05 was considered statistically significant).
W2-W6= Weeks of Imiquimod treatment, LT=Littermate, MLN= mesenteric lymph nodes, DCs= Dendritic Cells.
