## Supplementary material for "TLR7-Dependent Autoimmunity Remodels Intestinal Immune Cell Dissemination": Table2

**Table S2. Complementary analysis of intestinal immune dissemination associated with the breeding environment**

| B Cells | Control | Cohoused | P Value | Significance |
|---|---|---|---|---|
| Blood | 86.910±3.695 | 89.70±2.359 | 0.9095 | ns |
| Spleen | 65.310±3.432 | 68.530±2.801 | 0.5601 | ns |
| Kidney | 54.820±3.053 | 62.060±4.848 | 0.1595 | ns |
| **CD4 T Cells** | **Control** | **Cohoused** | **P Value** | **Significance** |
| Blood | 0.984±0.391 | 0.326±0.117 | 0.2174 | ns |
| Spleen | 16.490±1.872 | 13.08±1.242 | 0.207 | ns |
| Kidney | 11.180±1.490 | 7.656±1.394 | 0.1729 | ns |
| **CD8 T Cells** | **Control** | **Cohoused** | **P Value** | **Significance** |
| Blood | 0.890±0.295 | 0.729±0.245 | 0.6454 | ns |
| Spleen | 7.748±1.003 | 5.704±0.603 | 0.1056 | ns |
| Kidney | 6.321±0.830 | 4.524±0.988 | 0.0765 | ns |
| **Monocytes** | **Control** | **Cohoused** | **P Value** | **Significance** |
| Blood | 0.000±0.000 | 0.000±0.000 | >0.9999 | ns |
| Spleen | 0.000±0.000 | 0.000±0.000 | >0.9999 | ns |
| Kidney | 0.000±0.000 | 0.000±0.000 | >0.9999 | ns |
| **Neutrophils** | **Control** | **Cohoused** | **P Value** | **Significance** |
| Blood | 0.442±0.226 | 0.728±0.291 | 0.4201 | ns |
| Spleen | 0.0346±0.013 | 0.080±0.022 | 0.3018 | ns |
| Kidney | 2.230±0.970 | 4.030±2.550 | 0.5893 | ns |
| **Dendritic Cells (MHCII$^+$ CD11c$^+$)** | **Control** | **Cohoused** | **P Value** | **Significance** |
| Blood | 0.552±0.165 | 0.596±0.231 | 0.7182 | ns |
| Spleen | 0.521±0.147 | 0.438±0.094 | 0.8035 | ns |
| Kidney | 3.132±0.721 | 1.463±0.616 | 0.0352 | * |
| **CD11b$^+$ DCs** | **Control** | **Cohoused** | **P Value** | **Significance** |
| Blood | 0.301±0.111 | 0.101±0.048 | 0.1656 | ns |
| Spleen | 0.146±0.431 | 0.090±0.022 | 0.6542 | ns |
| Kidney | 1.262±0.370 | 0.000±0.000 | 0.0005 | *** |

Values represent frequencies (% of KikGR Red$^+$ cells) and are expressed as mean±SEM (n=9-18 mice per group).
Mann-Whitney test was performed and a P<0.05 was considered statistically significant.
*P<0.05; **P<0.01; ***P<0.001; ****P<0.0001; ns= not significant.
MLN= mesenteric lymph nodes, DCs= dendritic cells
