## Supplementary material for "TLR7-Dependent Autoimmunity Remodels Intestinal Immune Cell Dissemination": Table3

| Table S3. Intestinal immune dissemination following fecal microbiota transfer | | | | |
|---|---|---|---|---|
| **B Cells** | **Control FMT** | **TLR7.1 Tg FMT** | **P Value** | **Significance** |
| MLN | 59.740±7.185 | 55.050±2.642 | 0.6623 | ns |
| Spleen | 68.840±5.305 | 73.100±1.994 | 0.6623 | ns |
| Kidney | 75.600±5.684 | 77.200±1.636 | 0.9719 | ns |
| **CD4 T Cells** | **Control FMT** | **TLR7.1 Tg FMT** | **P Value** | **Significance** |
| MLN | 16.180±3.578 | 19.800±1.690 | 0.2468 | ns |
| Spleen | 15.270±2.684 | 14.220±1.253 | 0.9307 | ns |
| Kidney | 6.166±1.166 | 5.253±0.817 | 0.6623 | ns |
| **CD8 T Cells** | **Control FMT** | **TLR7.1 Tg FMT** | **P Value** | **Significance** |
| MLN | 10.950±1.455 | 12.410±0.758 | 0.303 | ns |
| Spleen | 7.336±1.544 | 5.870±0.409 | 0.5368 | ns |
| Kidney | 1.562±0.764 | 1.240±0.548 | 0.2294 | ns |
| **Monocytes** | **Control FMT** | **TLR7.1 Tg FMT** | **P Value** | **Significance** |
| MLN | 0.000±0.000 | 0.000±0.000 | >0.9999 | ns |
| Spleen | 0.000±0.000 | 0.000±0.000 | >0.9999 | ns |
| Kidney | 0.000±0.000 | 0.000±0.000 | >0.9999 | ns |
| **Neutrophils** | **Control FMT** | **TLR7.1 Tg FMT** | **P Value** | **Significance** |
| MLN | 0.000±0.000 | 0.000±0.000 | >0.9999 | ns |
| Spleen | 0.201±0.102 | 0.071±0.038 | 0.1255 | ns |
| Kidney | 0.0480±0.048 | 0.262±0.169 | 0.6667 | ns |
| **Dendritic Cells ($MHCII^+$ $CD11c^+$)** | **Control FMT** | **TLR7.1 Tg FMT** | **P Value** | **Significance** |
| MLN | 5.942±3.323 | 5.197±0.986 | 0.5368 | ns |
| Spleen | 1.172±0.399 | 0.635±0.051 | 0.1948 | ns |
| Kidney | 3.532±1.515 | 1.848±0.432 | 0.6623 | ns |
| **$CD11b^+$ DCs** | **Control FMT** | **TLR7.1 Tg FMT** | **P Value** | **Significance** |
| MLN | 0.378±0.131 | 0.198±0.042 | 0.5758 | ns |
| Spleen | 0.261±0.165 | 0.057±0.014 | 0.0866 | ns |
| Kidney | 2.546±0.998 | 1.110±0.270 | 0.4286 | ns |

Values represent frequencies (% of KikGR $Red^+$ cells) and are expressed as mean±SEM (n=5-6 mice per group).
Mann-Whitney test was performed and a P<0.05 was considered statistically significant.
*P<0.05; **P<0.01; ***P<0.001; ****P<0.0001; ns= not significant.
FMT= fecal microbiota transfer, MLN= mesenteric lymph nodes, DCs= dendritic cells
