## Supplementary material for "TLR7-Dependent Autoimmunity Remodels Intestinal Immune Cell Dissemination": Table4

**Table S4. Intestinal immune dissemination following short-term pathobiont colonization**

| **B Cells** | **Control** | ***E. faecalis*** | ***E. gallinarum*** |
|---|---|---|---|
| MLN | 49.160±7.295 | 38.190±8.649 | 53.310±3.973 |
| Spleen | 61.490±5.127 | 51.080±9.328 | 64.710±3.341 |
| Kidney | 59.260±4.800 | 41.110±9.832 | 49.080±5.562 |
| **CD4 T Cells** | **Control** | ***E. faecalis*** | ***E. gallinarum*** |
| MLN | 11.530±2.533 | 16.870±1.548 | 12.470±1.721 |
| Spleen | 17.970±2.827 | 23.870±4.781 | 15.370±1.743 |
| Kidney | 9.973±1.162 | 12.090±3.088 | 8.204±1.344 |
| **CD8 T Cells** | **Control** | ***E. faecalis*** | ***E. gallinarum*** |
| MLN | 11.070±2.109 | 9.528±0.641 | 9.744±1.120 |
| Spleen | 6.736±0.847 | 6.847±1.394 | 5.670±0.535 |
| Kidney | 4.630±0.977 | 4.280±1.740 | 5.526±0.950 |
| **Monocytes** | **Control** | ***E. faecalis*** | ***E. gallinarum*** |
| MLN | 0.002±0.002 | 0.005±0.005 | 0.002±0.001 |
| Spleen | 0.000±0.000 | 0.481±0.025[a] | 0.000±0.000 |
| Kidney | 0.369±0.162 | 0.370±0.278 | 0.187±0.075 |
| **Neutrophils** | **Control** | ***E. faecalis*** | ***E. gallinarum*** |
| MLN | 0.004±0.002 | 0.009±0.003 | 0.000±0.000 |
| Spleen | 0.124±0.030 | 0.206±0.068 | 0.226±0.072 |
| Kidney | 2.001±0.669 | 1.753±1.298 | 3.438±0.945 |
| **Dendritic Cells ($MHCII^+$ $CD11c^+$)** | **Control** | ***E. faecalis*** | ***E. gallinarum*** |
| MLN | 6.783±1.937 | 8.447±2.412 | 5.419±0.887 |
| Spleen | 0.558±0.106 | 0.843±0.197 | 0.915±0.150 |
| Kidney | 2.889±0.707 | 1.960±1.184 | 5.646±0.710[a] |
| **$CD11b^+$ DCs** | **Control** | ***E. faecalis*** | ***E. gallinarum*** |
| MLN | 0.619±0.161 | 2.248±0.922 | 1.190±0.253 |
| Spleen | 0.105±0.031 | 0.380±0.091 | 0.102±0.016 |
| Kidney | 0.571±0.142 | 0.830±0.457 | 0.967±0.235 |

Values represent frequencies (% of KikGR $Red^+$ cells) and are expressed as mean±SEM (n=6-14 mice per group).

[a]= significantly different from control group (Kruskal-Wallis test followed by Dunn's multiple-comparison test and a $P<0.05$ was considered statistically significant).

MLN= mesenteric lymph nodes, DCs= Dendritic Cells
